# Perturbation-guided mapping of colorectal cancer cell states to causal mechanisms

**DOI:** 10.64898/2026.03.03.708171

**Authors:** Soroor Hediyeh-zadeh, Tzen S. Toh, Olli Dufva, Giuseppe Serra, Rashika Jakhmola, Camille Fourneaux, Goncalo Rei Pinto, Zijian Fang, Gabriele Picco, Amanda J. Oliver, Rasa Elmentaite, Till Richter, Ken To, J. Patrick Pett, Sarah A. Teichmann, Elham Azizi, Florian Buettner, Fabian J. Theis, Mathew J. Garnett

## Abstract

Colorectal cancer (CRC) cell atlases have refined descriptive maps of tumour ecosystems, yet cross-sample integration often obscures disease-relevant patient-specific variation and remains largely correlative, limiting insight into the mechanisms and state transitions that drive progression and treatment response. Here, we develop a continual learning framework to construct a comparative single-cell CRC atlas spanning over 300 patients and 1.5 million cells, preserving inter-patient variation while aligning healthy and malignant contexts. We resolve distinct non-canonical malignant cell states, including an endoderm-like state enriched in microsatellite-stable, *KRAS*-mutant CRC with features of oncofetal plasticity. Cell states are recapitulated in patient-derived organoids, establishing a tractable model of reprogramming. By linking the observational atlas to a large-scale perturbation atlas using relative representations, we map perturbations that drive cells toward defined phenotypic extremes. We connect cell states to therapeutic responses, showing that MAPK inhibition induces a shift away from a proliferative phenotype and converges towards a plastic, endoderm-like state. Together, this framework moves beyond static atlases to enable mechanistic modeling of cell-state regulation and causal inference toward cell-state–directed therapies.

## Introduction

Cell atlases of CRC have characterized cellular variation across the available phenotypic landscape of CRC^1–11^. However, they are limited in their potential to impart an understanding of the mechanisms that govern cellular transitions across this landscape. Inferring these causal mechanisms is enabled by perturbation data and linking both observational and perturbation data remains a fundamental challenge for modeling cellular transitions^12^.

Malignant cells may co-opt multiple conserved or distinct states and exhibit cellular state transitions in response to therapy^13–15^ and disease progression in CRC^11^. Resolving these states and their transitions across disease contexts such as normal, pre-malignant, primary tumour, metastatic, and perturbation-induced states is required to identify axes of therapeutically actionable transitions. This also requires careful integration of cancer single-cell RNA-sequencing (scRNA-seq) data that accounts for patient-specific variation^15,16^. However, current methods for scRNA-seq data integration often over-correct tumour-specific variation, thereby obscuring disease-relevant states. A method that preserves both shared healthy programmes and patient-specific disease variation could enable comparison of cells across malignant contexts and facilitate integration of perturbation data^17–19^.

Mapping perturbation cell atlases onto an observational phenotypic space enables identification of perturbation-induced transition axes and links cell-state changes to mechanism, with potential to specify perturbations driving therapeutic vulnerabilities or interventions. Pre-clinical models of CRC have been used to model therapeutic responses to small molecule perturbations at scale and identify potential therapeutic targets^20–25^. Their full potential remains unrealized for use in understanding how CRC cells undergo cellular state changes during adaptation to therapy^13,14,26^ as they should be contextualized within a detailed map of the CRC phenotypic landscape with high confidence^27^. Here, we introduce a comparative cell atlas, designed to balance alignment of shared cell states with retention of context-specific variation during data integration, thereby providing an enhanced contextual map to study CRC. We construct a comparative CRC atlas spanning normal, pre-malignant, primary tumour, and metastatic states while retaining patient-specific variation, and link it with the Tahoe-100M perturbation dataset^28^ in a shared phenotypic space to resolve perturbation-induced malignant state transitions and their mechanisms.

## Results

### Constructing a comparative single cell CRC atlas using continual learning

Implementing a comparative atlas across heterogeneous cancer datasets requires a framework that incrementally incorporates new case-control data while preserving previously learned biological structures. We developed an incremental atlas expansion strategy based on continual learning (CL) that updates an existing reference atlas with case-control scRNAseq query data by aligning the healthy reference and query-control cells. A two-part specialized regularization technique (**Fig. 1a, Methods**) increases the adaptability of the conditional Variational Autoencoder (cVAE) model, commonly used for removing technical effects and constructing integrated reference cell atlases, to new variation as additional datasets are added, while retaining old variation (**Methods**). The incremental update with regularization encourages preservation of both healthy and disease-specific states enabling comparative analysis of cellular states (**Supplementary Note 1-3**). Using published scRNA-seq case-control CRC datasets (query) and an existing gut atlas^29^, we built a comparative CRC atlas in a single incremental step (**Fig. 1b**).

**Fig. 1:**
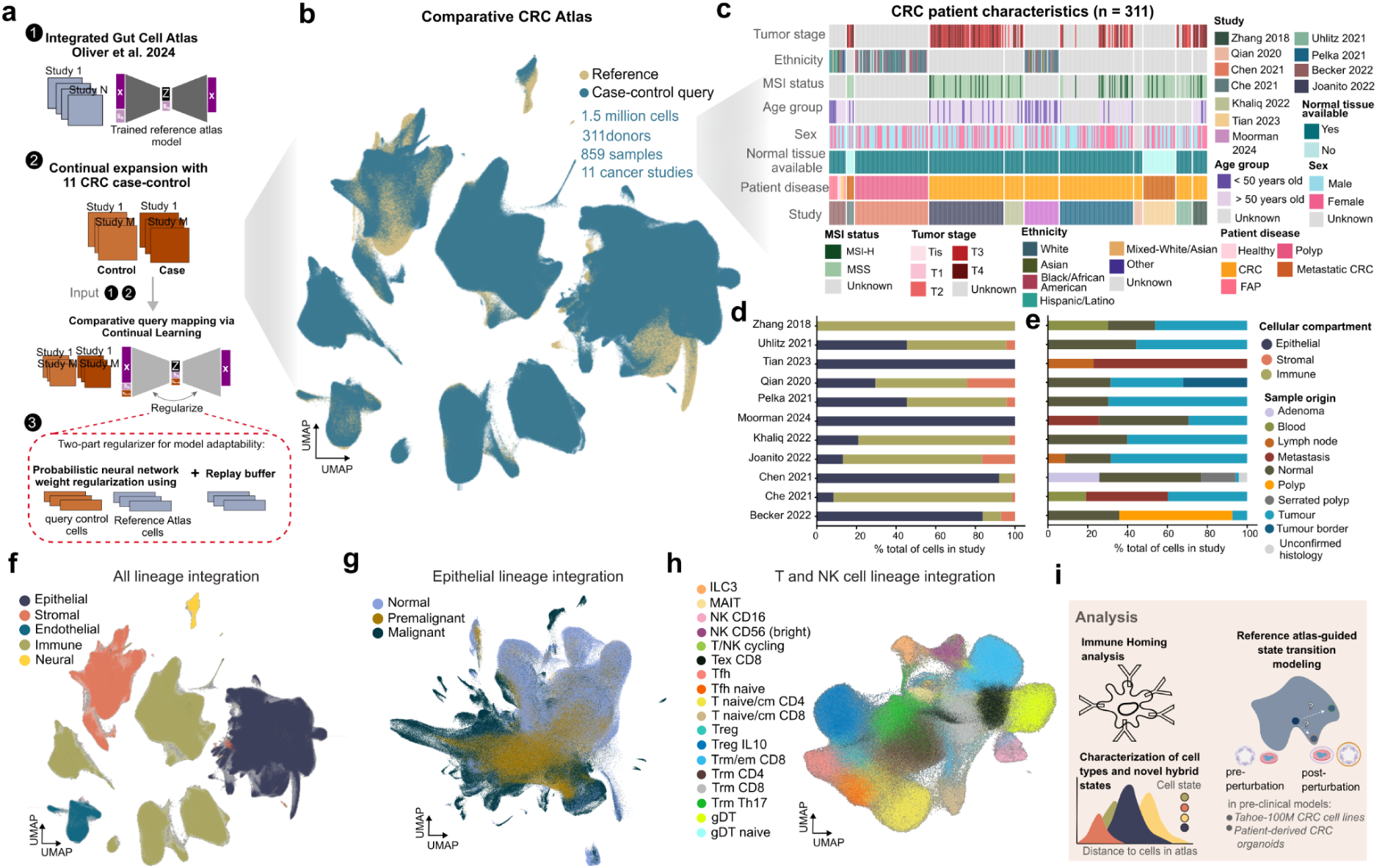
Comparative CRC atlas construction. **(a)** Schematic of our comparative atlas construction workflow. We built a comparative atlas by (1) incrementally updating an existing reference gut epithelial cell atlas, with (2) query case-control datasets. Rather than freezing model weights, the model is continually trained on the query case-control data. A regularizer penalizes updates to the neural network weights based on their importance for reconstruction of the states in the reference and query-control cells and a Replay buffer (**Methods**) (3). **(b)** A comparative CRC atlas constructed by expanding the integrated gut atlas (Oliver *et al*.^29^) with colorectal cancer cell datasets in a single incremental step. **(c)** An overview of patient characteristics, **(d)** cellular compartments, and **(e)** sample origin by dataset. **(f-h)** UMAP embedding of all lineage, epithelial lineage, and T and NK cell lineage integration respectively. **(i)** Schematic of comparative CRC atlas applications used here.

We selected and curated eleven publicly available high quality CRC scRNA-seq datasets^1–11^ consisting of over 1.5 million cells, 311 donors, and 859 samples (**Fig. 1c, Supplementary Table 1**). Nine of these datasets had healthy cells and so were suitable for our CL approach. These datasets had donor representation across a variety of ages, ethnicities, sex, and clinical metadata such as microsatellite instability (MSI) status, tumour stage, and patient disease information which allowed the later interrogation here of cellular variation with clinical co-variates. All of the datasets contained epithelial cells across different disease contexts apart from Zhang *et al*.^9^ which contained only immune cells (**Fig. 1d**). These cells were profiled from various samples including those from normal tissues, polyps, primary tumours, and metastatic tumours (**Fig. 1e**).

The tumour microenvironment (TME) of CRC has been a primary area of focus in most published studies, including integrative analysis of CRC^30^. A systematic integration of malignant epithelial cells in CRC remains lacking as current methods for scRNA-seq data integration focus on resolving cellular variation but do not fully account for patient-specific variation, diluting meaningful biological signals^31^. We therefore focus primarily on cell state changes in epithelial cells across the malignant continuum of CRC as our CL approach is designed to preserve patient-specific variation (**Methods**).

We built the comparative CRC atlas by performing data integration at three levels; an all cell type lineage integration (**Fig. 1f**), an epithelial cell integration (**Fig. 1g**), and a T and natural killer (NK) cell lineage integration (**Fig. 1h**). We include the integration of the T and NK cells to study malignant cell-immune cell interactions given their importance in influencing CRC patient responses to immunotherapies^32,33^. We used harmonized, author-provided cell type annotations for the all cell type lineage integration to initially test the performance of the CL approach. As expected, after data integration, cells were primarily grouped by the five main cell types lineages (**Fig. 1f**). We proceeded to primarily focus on integrations of epithelial cells (n = 800,350 cells, **Fig. 1g**) and T and NK cells (n = 439,924, **Fig. 1h**). **Fig. 1i** summarizes some of the analysis enabled by the comparative CRC atlas.

### The comparative CRC atlas resolves tumour cell populations across disease contexts

We term the final comparative epithelial cell atlas the Epi-CRC atlas. We observed that the epithelial cell integration preserved expected normal colonic cell types, with query-control cells (normal cells from the CRC datasets) aligning closely to the reference. In contrast, malignant cells, including those from metastatic samples, exhibited pronounced patient-specific variation, forming distinct clusters with low mixing in the embedding (**Fig. 2a, Extended Data Fig. 1a,d**).

**Fig. 2:**
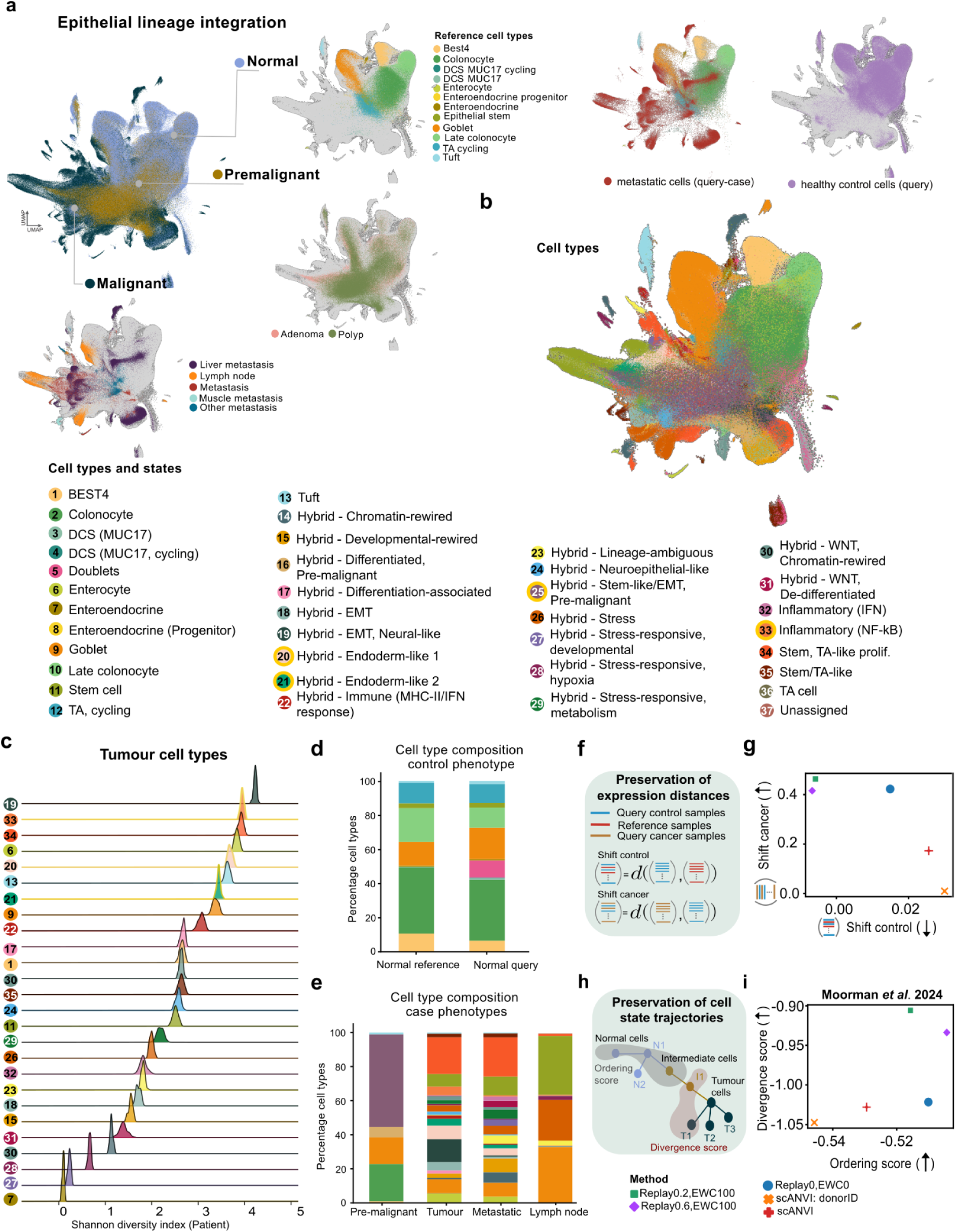
Continual learning improves model adaptability and recapitulates malignant progression in CRC. **(a)** The Epi-CRC atlas captures the normal to malignant continuum through pre-malignacy. **(b)** UMAP embedding of cells coloured by cell types. The states discussed in more detail identified from Epi-CRC integration are highlighted with a gold outline. **(c)** Shannon Diversity Index for tumour cell types. Diversity was measured with respect to patient IDs to identify shared malignant states across the comparative atlas. **(d)** Cell type composition in normal samples and **(e)** across tumour contexts. **(f)** Schematic of shift distance metrics. **(g)** Distance of tumour samples to normal samples in the query (larger values indicate better performance) versus distance between healthy samples in the reference and query-control samples (smaller values indicate better performance). Models in the top-left corner perform best in preserving shared healthy and novel disease states indicating high adaptability (**Supplementary Note 2**). **(h)** Schematic of the Ordering Score and Divergence Score used to assess integration. **(i)** Divergence and ordering score for capturing transition to malignancy at the cell level using cells from Moorman *et al*.^11^ as reference in the Epi-CRC atlas only. Models in the top-right corner perform the best in capturing cell trajectories.

We clustered the embedding of the diseased cells after data integration and annotated the clusters using established marker genes where possible (**Methods**)^1,34^. However, for the majority of malignant epithelial cell clusters, the expression of cell type markers were non-uniform (**Extended Data Fig. 2a**) which prompted us to also investigate transcription factor (TF) activity across the diseased epithelial cells. We hypothesized that the diseased epithelial cells retained some features of normal cells, but had TF profiles that distinguished them from each other. By using both marker gene expression and TF activity scoring for the malignant cells (**Extended Data Fig. 2b**), we annotated 37 distinct epithelial cell types and states across the Epi-CRC atlas (**Fig. 2b**). We termed many of these states “hybrid” (n = 20), given that they did not fit canonical cell type markers from the healthy colon or CRC (**Extended Data Fig. 2a,c, Methods, Supplementary Table 2**).

We calculated the Shannon diversity index (SDI) — a measure of diversity in a population — for the 26 malignant cell types and states with respect to patients to exclude confounding study-specific variation. We found 9 states exhibited a high SDI (SDI larger than 3, range 0.09-4.22, **Fig. 2c**) suggesting that they are highly conserved across patients, and 2 states showed a low SDI (less than 0.5) suggesting that they exist in a limited number of patients. Normal cells in the query retained compositional similarity to normal cells in the reference (**Fig. 2d**). Pre-malignant epithelial cells (n = 157,467), from samples both matched and unmatched to primary tumours, consisted of a mix of stem-like and differentiated cell types (**Fig. 2e**). A hybrid, stem-like/EMT state with high activity of TFs involved in EMT, stemness, and embryonic development (*SOX4*, *SOX9*, *DACH1* and *SNAI2*) formed 54% of epithelial cells in the pre-malignant samples (cell state 25; **Fig. 2e**), suggesting it is a key feature of pre-malignant transition towards malignancy^3,5^.

Overall, cell type and state proportions were similar for primary tumours and metastatic samples (**Fig. 2e**). The dominant cell state across both was a stem, TA-like proliferative state (state 34) consistent with the highly proliferative nature of tumours. Stem cell (state 11) proportions were higher in the primary tumour population (7.4%) compared to the metastatic tumours (4.9%). A notable exception was the relatively high proportion of stem cells in samples from lymph nodes, suggesting potential infiltration of cells from the tumour^35,36^ (**Fig 2e**). Collectively, our integration with CL conserves the continuum from normal to malignant states, identifies recurrent non-canonical hybrid cell states, and allows resolution of tumour cell populations across disease contexts.

### Epi-CRC atlas preserves features of disease progression

Architectural surgery refers to the transfer of learned model weights from a reference dataset to map new query datasets, and is an established transfer learning approach for extending existing reference atlases^37^. Rather than simply transferring model weights, our CL framework enables iterative weight updates as the reference model is continually trained on new query datasets. We use a subset of query-control together with healthy reference cells to guide the direction of the updates so that previously captured variation in the reference model is preserved.

This weight regularization is implemented using Elastic Weight Consolidation (EWC)^38^, which we modified to accommodate case-control data (**Methods**), thereby aligning control cells from both the reference and incoming datasets. In a separate step, a representative subset of reference cells are selected and *replayed* (rehearsed) during model updating to maintain the global structure of the reference atlas during incremental expansion. Together, this two-step regularization strategy improves model adaptability while preserving prior knowledge, enabling incremental updates to the atlas (**Fig. 1a, Methods, Supplementary Note 1-2**).

We used two metrics, one based on expression distances (**Fig. 2f**) and one based on disease trajectories (**Fig. 2h**), to compare the performance of our CL method versus architecture surgery (**Supplementary Note 1**). First, we compared distances between query-control and cancerous samples in our CRC dataset (shift-cancer), and between query-control and healthy samples in the reference from Oliver *et al*.^29^ (shift-control) (**Fig. 2f, Methods**). While shift-control measures how well the control and normal samples from query and reference are integrated, shift-cancer measures preservation of tumour-specific variation after integration. Using our continual learning (CL) framework with regularization parameters (replay = 0.2, EWC = 100), tumour samples showed greater separation from control samples while preserving minimal distance between reference and query-control samples, compared to integration via architecture surgery (**Fig. 2g**). Second, to evaluate whether our CL approach recapitulates progression from normal to metastatic disease, we used samples from Moorman *et al.*^11^, which provides a well-characterised reference for this transition. We investigated whether any two adjacent cell or disease states on the known trajectory are adjacent neighbors on the integrated embedding (the ordering score), and if any two non-adjacent states sharing the same origin on the trajectory are non-adjacent neighbors (the divergence score)^17^ (**Fig. 2h, Methods**). The ordering score measures preservation of cell state orders on the trajectory whereas the divergence score measures preservation of branching points on the trajectory. We found embeddings from our CL approach – (Replay 0.2,EWC100) and (Replay 0.6,EWC100) – had both higher ordering and divergence scores compared to the two scANVI (Transfer Learning) baselines (**Fig. 2i, Supplementary Note 2**).

We also found that SDI is smaller for malignant cells compared to cells from normal or pre-malignant stages (**Extended Data Fig. 1d**), suggesting that the integrated embedding is capturing shared healthy states and more heterogeneous malignant states. Additionally, sample-level embeddings from healthy individuals in the reference and query-controls were better aligned on the comparative mapping embedding compared to architecture surgery (**Extended Data Fig. 1e**) suggesting that sample trajectories are better preserved by the CL regularization approach.

We also benchmarked our CL regularization scheme against *de novo* integration and architecture surgery using simulated PBMC datasets, modeling disease-specific transcriptional shifts and cell type compositional changes (**Supplementary Note 3**). We evaluated the different parameters for improving integration performance and cVAE model adaptability (**Supplementary Note 3**).

We show our CL scheme on real and simulated data effectively balances removal of technical variation with preservation of biological variation while maintaining expected disease trajectory. The confidence in our approach allows the identification of hybrid malignant cell states which do not recapitulate marker gene expression of healthy colonic cells. It enables a more complete annotation of the Epi-CRC atlas and can be used to interrogate relationships between malignant cell type and states with CRC genetics, immune cell composition, and clinical data.

### Endoderm-like malignant states are associated with microsatellite stability, *KRAS* mutation, and immune exhaustion

Leveraging the size and detailed curation of the fully annotated Epi-CRC atlas (**Fig. 2b**), we sought to investigate disease-relevant cell types and states and their relationship to tumour genetics, immune cell composition, and clinical co-variates. We nominated the 10 cell types and states with the highest SDI (**Fig. 2c**) for this analysis given they are conserved across patient populations.

We investigated cellular composition across microsatellite stable (MSS) and microsatellite Instability-High (MSI-H) CRCs given their varying prognosis by MSI status^33,39^ and revealed differences in their cell type enrichment. In particular, MSS CRCs were enriched for the hybrid - endoderm-like 1 state (cell state 20, two-sided Fischer’s exact test odds ratio = 9.5 adjusted p-value < 0.05) and hybrid - endoderm-like 2 state (cell state 21, two-sided Fischer’s exact test odds ratio = 106.6 adjusted p-value < 0.05), while an inflammatory (NF-κB) state (cell state 33) was enriched in MSI-H CRCs (two-sided Fischer’s exact test odds ratio = 0.105, adjusted p-value < 0.05, **Fig. 3a**).

**Fig. 3:**
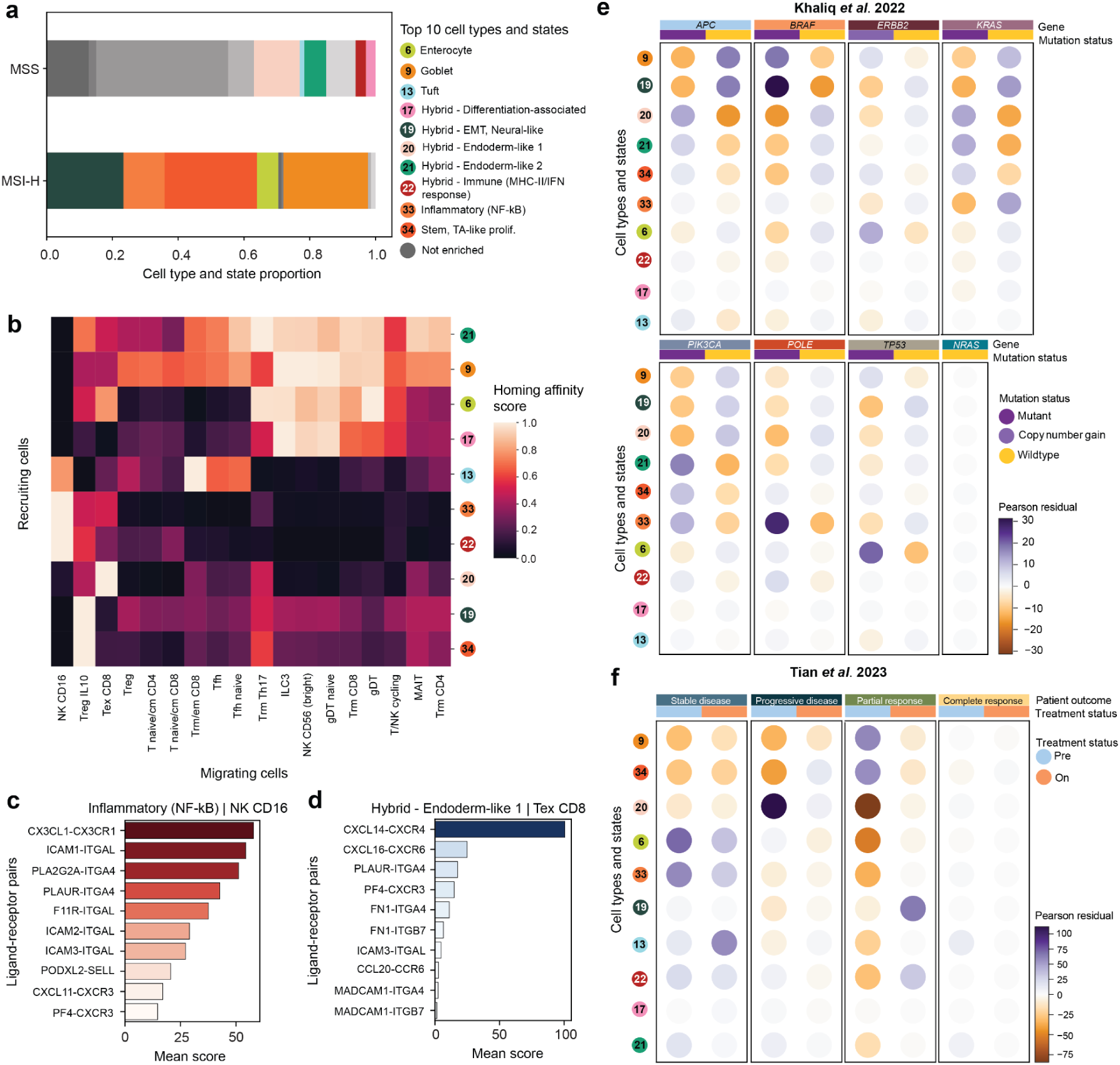
Endoderm-like states are enriched in MSS CRC and are associated with *KRAS* mutation. **(a)** Differential cell types and states in MSS versus MSI-H CRC. Enriched cell types and states in each are coloured (two-sided Fischer’s exact test, adjusted p-value < 0.05) and not enriched are greyed. **(b)** The homing affinity score between epithelial cell types with the top 10 highest Shannon Diversity Index values and migrating immune cells. A higher score reflects greater affinity. **(c)** Top 10 ranked predicted ligand-receptor pairs between natural killer CD16 cells and inflammatory (NF-κB) state. **(d)** Similar to **(c)** but for exhausted CD8 T-cells and the hybrid - endoderm-like 1 state. **(e)** Relationship between cell types and states with mutation status from cells from Khaliq *et al*^2^. Positive Pearson residuals indicate a positive association between the cell types and states with mutation status (purple) and vice versa for negative Pearson residuals (orange). **(f)** Similar to **(e)** but between cell states and pre-treatment (left) and on-treatment (right) patient outcomes from Tian *et al*^6^.

The hybrid - endoderm-like 1 state (cell state 20) is defined by high activity for TFs involved in developmental patterning (*HAND1*, *ISL1*, *NR6A1*), BMP signaling (*SMAD1*), and hepatocyte identity (*HNF1A*, *FOXA3*) (**Extended Data Fig. 2b, Supplementary Table 2**). While *HNF1A* and *FOXA3* are associated with hepatocyte cell fates, they are also expressed in organs of endoderm origin^40,41^. Differential gene expression analysis between all cell types and states in the Epi-CRC atlas revealed that the hybrid - endoderm-like 1 state (cell state 20) had genes involved in WNT-signalling (*ASCL2*) and EGFR signaling (*EREG, AREG*) differentially expressed (**Supplementary Table 3**). In contrast, the hybrid - endoderm-like 2 state (cell state 21) had high TF activity for TFs involved in WNT signaling (*TCF4)* and early development *(GATA6, HNF4A, FOXI1)* (**Extended Data Fig. 2b, Supplementary Table 2**). It had a mix of developmental related genes (*SOX4*, *MYC*, *PROX1*) and WNT-signalling (*ASCL2*) differentially expressed. *MEX3A*, a marker of drug-tolerant persister cells in CRC^42^ was also among differentially expressed genes in the hybrid - endoderm-like 2 cell state (cell state 21) when compared between all cell types and states in the atlas (**Supplementary Table 3**). The TF activity and differentially expressed genes indicate converging but distinct malignant programmes between these two states.

CRC patients with MSI-H tumours have high tumour mutational burdens and exhibit better responses to immune checkpoint inhibition^33,39^. We hypothesized that these cell states from the Epi-CRC atlas may exhibit differences in their interactions with their immune cell counterparts given that CRCs have variation between immune cell infiltration which contributes to variation in clinical outcomes^1,33,39^. We calculated an affinity score by multiplying a signature based on chemoattractant ligand expression (source) from the epithelial cells with the receptor (target) gene expression of the immune cells (**Methods**). A high affinity score represents a high probability of interaction between two groups of cells. We found that tissue-resident memory T cells (Trm), type 3 innate lymphoid cells (ILC3), γδ T cells (gdT), and CD56bright NK cells had a high affinity score towards mostly normal cell types. Tumour epithelial cell types show a high affinity score towards exhausted T cells (Tex), CD16+ NK cells, and IL10+ Tregs. Among tumour epithelial cells, inflammatory (NF-κB) and hybrid - immune (MHC-II/IFN response) cell types were predicted to interact with CD16+ NK cells while the hybrid endoderm-like 1 state attracted Tex (**Fig. 3b**).

We examined specific receptor–ligand interactions associated with the inflammatory (NF-κB) state (state 33) that are predicted to promote CD16⁺ NK cell migration. Among these, the *CX3CL1–CX3CR1* axis emerged as the top predicted interaction linking inflammatory tumour cells to CD16⁺ NK cells. Genes involved in cell adhesion (*ITGA4* and *ITGAL*) were also among the top 10 ligands found in this interaction (**Fig. 3c**).The *CX3CL1-CX3CR1* axis is associated with immune cell recruitment and better CRC patient outcomes^43^, suggesting the inflammatory (NF-κB) state promotes a favourable immune microenvironment. We next characterised receptor–ligand interactions between the hybrid endoderm-like 1 state (state 20) and exhausted Tex cells, identifying *CXCL14–CXCR4* as the top predicted interaction mediating this crosstalk (**Fig. 3d**). The role of *CXCL14* in CRC has been more extensively reported in cancer-associated fibroblasts (CAFs)^1,44^ and has been previously reported to be associated with poorer prognosis in patients with stage III/IV disease^45^ thus suggesting that the hybrid - endoderm-like 1 state may contribute to an immunosuppressive microenvironment.

Using patients from Khaliq *et al*.^2^, we identified the hybrid - EMT, neural-like state (state 19) to be the most positively associated with *BRAF*-mutant CRC, while the inflammatory (NF-κB) state (state 33) was positively associated with *POLE*-mutant CRC in patients with available cancer driver mutation annotation. Both endoderm-like states were positively associated with *KRAS*-mutant CRC (Chi-squared = 7,878.2, degrees of freedom = 126, p-value < 0.05, **Fig. 3e**).

The enrichment of the endoderm-like cell states with MSS CRC and their positive association with *KRAS* mutation in our analysis are reminiscent of oncofetal states in KRAS inhibitor resistant *KRAS*-mutant CRC^13^ and that drive metastatic progression^11^. We investigated the presence of oncofetal states in the endoderm-like states identified from the Epi-CRC atlas. We found the hybrid - endoderm-like 1 state (state 20) had high *CLU* and *EMP1* expression suggesting an invasive phenotype^46^. Compared to the stem cells from malignant samples, the hybrid - endoderm-like 2 state (state 21) exhibited high expression of YAP/TEAD genes, low intestinal stem cell (ISC) marker gene (*LGR5, SMOC2*) expression, and elevated expression of endoderm development (*PROX1, NKD1*) genes (**Extended Data Fig. 3a**). We calculated proliferative cancer stem cell (proCSC) and revival CSC (revCSC) signatures^47^ in the endoderm-like states compared to other stem cell populations in the comparative CRC atlas to further characterize these features (**Extended Data Fig. 3b and 3c**). The hybrid - endoderm-like 2 state was the highest scoring for the revCSC signature. Together, these results corroborate the hybrid - endoderm-like 2 state as slow cycling with features of a plastic, fetal-like transcriptional programme linked with therapy resistance and advanced disease^11,13^.

Our Epi-CRC atlas includes patient outcome data from Tian *et al.*^6^, who evaluated combined PD-1, BRAF, and MEK inhibition in *BRAF*-mutant CRC, enabling us to examine the relationship between cellular phenotypes and treatment outcome. Enterocytes (cell type 6), tuft cells (cell type 13), and the inflammatory (NF-κB) state (cell state 33) were associated with stable disease for patients on treatment. During treatment, the stem, TA-like proliferative cell state (state 34) had a weak positive association with progressive disease but pre-treatment had a negative association with progressive disease, suggesting that proliferative cell types are a feature of on-treatment resistant tumours. Additionally, the hybrid - EMT, neural-like and hybrid - immune (MHC-II/IFN response) states (state 19 and 22 respectively) were associated with partial response on-treatment. The hybrid - endoderm-like 1 state (state 20) was the only state with a positive association with progressive disease pre-treatment and during treatment (Chi-squared = 36,919, degrees of freedom = 63, p-value < 0.05, **Fig. 3f**).

Collectively, we found that endoderm-like cell states are characterized by features of early development and are of a slow-cycling nature, they are enriched in MSS CRCs and are associated with *KRAS* mutation, immune exhaustion, and poorer treatment responses. In contrast, MSI-H tumours were found to be enriched for the inflammatory (NF-κB) state and had high affinity for cytotoxic immune cells^33^. Moreover, the Epi-CRC atlas links cell states with clinical features to provide an integrative view of disease progression and clinical outcomes.

### Relative representations links observational tumour states to perturbation-induced transitions

Perturbation and observational cell atlases are typically analysed in isolation, and frameworks to map perturbation-induced transitions onto phenotypic landscapes defined by observational atlases remain limited. Establishing this link would connect defined cell states to the mechanisms that specify them and provide a principled strategy to identify therapeutic interventions that modulate or target specific cellular states. We used relative representations^48^ to link our observational Epi-CRC atlas as a reference to a perturbation atlas, the Tahoe-100M scRNA-seq dataset^28^, subsetted to 9 CRC cell lines and 804,879 cells perturbed with 380 small molecules (**Fig. 4a, Methods**). We also performed analysis on 3 Sanger CRC organoids profiled in house with scRNA-seq (**Methods**) in addition to publically available scRNA-seq data of matched organoids derived from a patient with primary (OKG146P) and metastatic (OKG146Li) CRC tumours^11^ to systematically compare the fidelity of different CRC pre-clinical models to tumour samples from patients for mechanistic modeling.

**Fig. 4:**
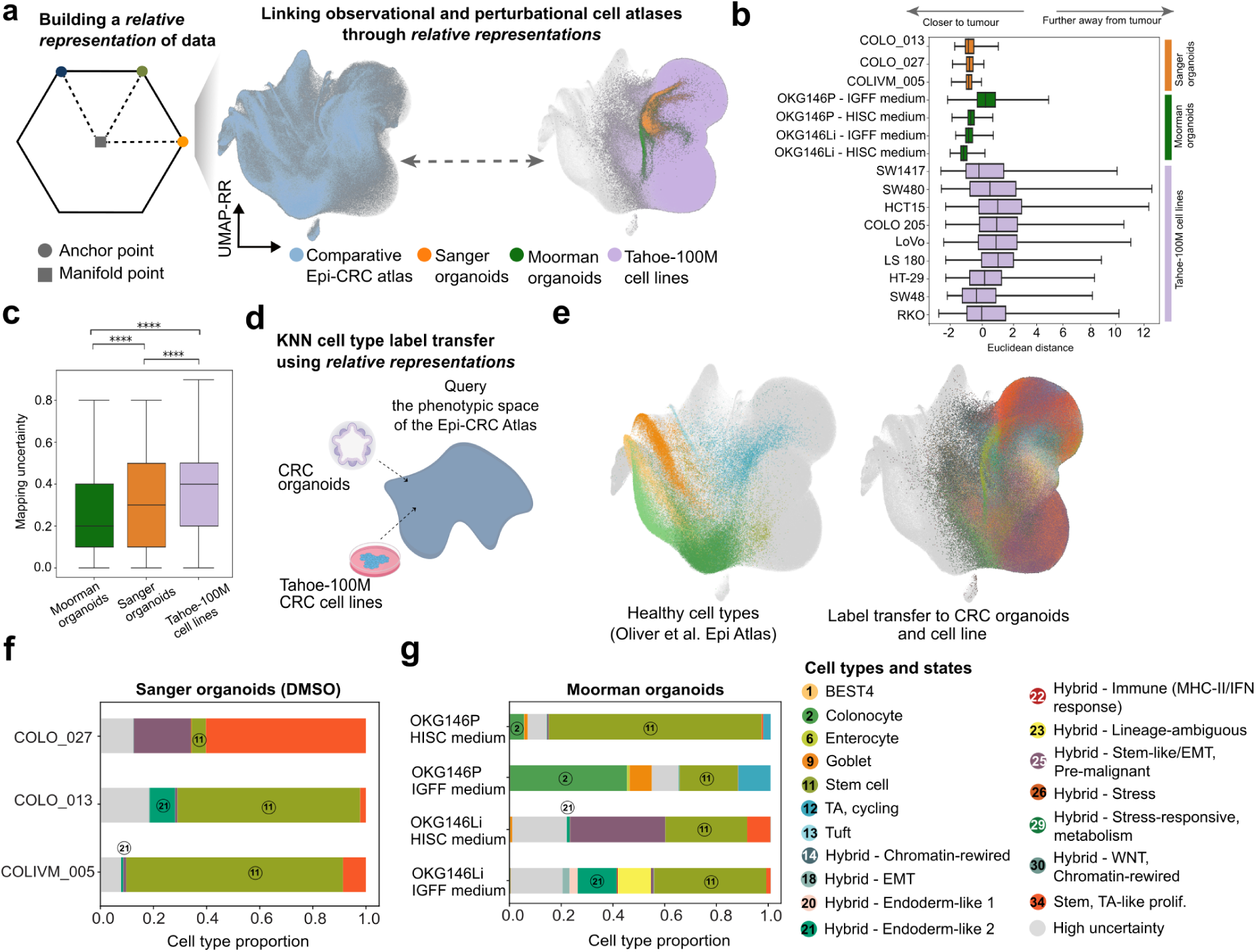
Relative representations for comparative analysis of cellular states in perturbational data reveal CRC organoids recapitulate important malignant states. **(a)** Schematic representation of relative representations (RR). UMAP projection of RRs showing cells from the observational Epi-CRC atlas, Tahoe-100M CRC perturbational atlas, and the organoids. The cells from the two atlases communicate in the space defined by RRs. **(b)** Median-normalised distance of single cells from pre-clinical models to tumour samples in the comparative Epi-CRC atlas. The smaller the distance, the closer to tumour samples. **(c)** Distribution of mapping uncertainty scores in CRC organoids and Tahoe-100M CRC cell lines. **(d)** Schematic of label transfer from the Epi-CRC atlas to CRC pre-clinical models on RR space**. (e)** UMAP projection of RR showing the original cell types in the reference from Oliver *et al*.^29^ (left) and the transferred cell types in the CRC organoids and Tahoe-100M cell lines (right). Grey cells in the left panel are cells that are not belonging to the reference while grey cells in the right panel are cells that are not belonging to CRC cell lines and organoids. **(f-g)** Proportion of predicted cell types and states in Sanger CRC organoids and Moorman CRC organoids, respectively. Cells with high uncertainty of assignment from label transfer are coloured grey. In the box plots in **b** and **c**, the center represents the median; bounds show the 25% and 75% percentiles; and whiskers indicate values within 1.5× the interquartile range.

We selected a set of cells across our observational atlas as anchors to cover the diversity and scale of the Epi-CRC atlas (**Fig. 4a**) and computed the similarity between the latent representations of these anchors and every cell in both atlases, independently for each latent dimension, to derive a new coordinate system in which every cell is represented relative to the anchor cells. In this space, cells from the observational and perturbational atlases are linked through relative representations, which we denote as RR hereafter (**Methods**).

We found all CRC organoids were closer towards the cell tumour samples in the comparative CRC atlas with the exception of the OKG146P organoid cultured in intestinal growth-factor-free (IGFF) medium compared to Tahoe-100M CRC cell lines (**Fig 4b**). The mapping uncertainty based on RR distances to the atlas was higher in the Tahoe-100M CRC cell lines (median = 0.34) compared to the Sanger CRC organoids (median = 0.28) and organoids from Moorman *et al*.^11^ (median = 0.25, Mann-Whitney-Wilcoxon test adjusted p-value < 0.05; **Fig. 4c**). This mapping uncertainty may underlie the greater dispersion of CRC cell line distances.

We performed *k*-NN label transfer to both the CRC organoids and cell lines to interrogate cell states from CRC tumours in both pre-clinical models (**Fig. 4d**). The structures of healthy cell types were preserved after transformation to new coordinates using relative representations (**Fig. 4e**, left panel) and the pre-clinical models were annotated with the 37 cell types and states from the Epi-CRC atlas (**Fig. 4e**, right panel). The Sanger CRC organoids consisted primarily of stem cells (cell type 11) and stem, TA-like proliferative cells, consistent with their in vitro culture conditions based on WNT-dependent stem cells from the colon^49^. The hybrid - endoderm-like 2 cells (state 21) were most abundant in COLO_013 (*KRAS-*mutant) followed by COLIVM_005 (*KRAS-*mutant) then COLO_027 (*BRAF-*mutant, **Fig. 4f**). This is concordant with our earlier observation that this state is positively associated with *KRAS* mutations (**Fig. 3d**).

The Moorman CRC organoids had cell state composition that differed by organoid and culture condition. OKG146P (primary tumour) organoids cultured in human intestinal stem cell (HISC) medium had colonocyte (cell type 2) and stem cells (cell type 11, **Fig. 4g**). OKG146Li (liver metastasis) organoids cultured in HISC medium had a small proportion of the hybrid - endoderm-like 2 state and higher levels of the hybrid - stem-like/EMT state from pre-malignant samples. Despite being cultured in IGFF medium, we found OKG146P retained largely similar composition but with higher proportion of colonocytes and OKG146Li had a higher proportion of the hybrid - endoderm-like 2 state (**Fig. 4g**). Our findings are consistent with previous observations that OKG146P retained higher levels of differentiated intestine phenotypes while OKG146Li had stem-like and endodermal states^11^.

Cells from the Tahoe-100M CRC cell lines were classified as high uncertainty (uncertainty score > 0.5) at a higher proportion (22%) compared to cells from the Sanger CRC organoids (17%) and Moorman CRC organoids (14%, **Extended Data Fig. 4a**). Most CRC cell lines were dominated by the hybrid - stem-like/EMT state associated with pre-malignant samples (state 25) and stem, TA-like proliferative states (state 34) while having a higher proportion of “high uncertainty” labeled cells (**Extended Data Fig. 4b and 5a**). EMT and cell cycle related gene expression programmes have been identified in previous systemic analysis of cancer cell lines^50^. CRC organoids had a lower proportion of cells assigned as “high uncertainty” (uncertainty score > 0.5, **Fig. 4f**, **Extended Data Fig. 5b and 5c**) in assigned cell types and states compared to the Tahoe-100M CRC cell lines (**Extended Data Fig. 4b**). Overall, our analysis suggests that organoids more closely represent cell types and states observed in patient tumours^11,51^ compared to cell lines. Moreover, our relative representation framework allows mapping of pre-clinical models to the Epi-CRC atlas to interrogate pre-clinical model cellular composition and further interrogation of cellular transitions induced by perturbations.

**Fig. 5:**
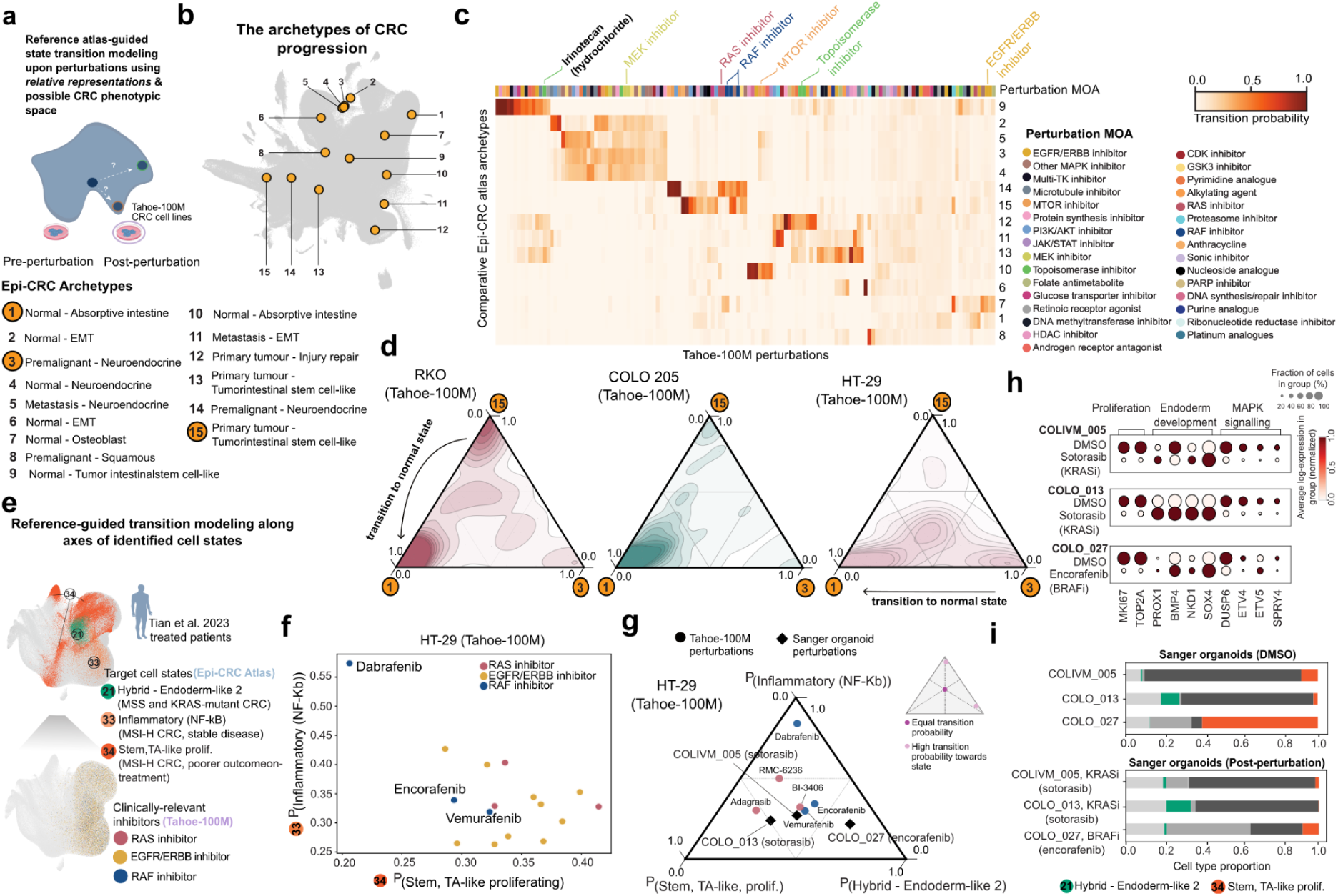
Reference-guided state transition modeling reveals perturbations that induce transitions along therapeutically-relevant cell state axes. **(a)** Schematic of our reference-guided framework for quantifying state transitions post perturbations using an observational cell atlas and RRs anchored against the target state neighbourhood. **(b)** Archetype analysis across epithelial cells identifies 15 extreme phenotypic states capturing CRC progression. **(c)** Quantitative analysis of cell state transitions induced by the Tahoe-100M perturbations towards Epi-CRC archetypes. Compounds are grouped by mechanism of action (MOA). **(d)** Quantifying cell state transitions induced by Tahoe-100M perturbations in three *BRAF*-mutant cell lines toward normal, pre-malignant, and malignant archetypes. **(e)** Schematic of framework for reference-guided transition modeling along axes of identified cell states. **(f)** Dabrafenib (BRAF inhibitor) induces transitions towards the inflammatory (NF-κB) state (associated with stable disease) in HT-29. **(g)** Distribution of the transition probability scores along axes of cell states induced by RAF and RAS inhibitors in HT-29 and three CRC organoid models. Circles represent RAS (red) and RAF (blue) inhibitors in Tahoe-100M. The Sanger organoids are marked as black diamonds. **(h)** Marker gene expression of proliferation, endoderm development, and MAPK signaling genes in the DMSO and treated Sanger organoids. **(i)** Cell type and state compositional changes in Sanger CRC organoid models upon treatment. The bars coloured in different shades of grey are the other remaining assigned cell types and states.

### Reference-guided perturbation modelling reveals context-dependent cell-state transitions

Having mapped perturbation responses onto the Epi-CRC atlas using relative representations, we examined how compounds from Tahoe-100M^28^ drive cell-state transitions to defined phenotypic states (**Fig. 5a**). We defined 15 phenotypic extremes (archetypes) spanning normal epithelium to malignancy (**Fig. 5b; Methods**) and computed relative representations to quantify shifts toward each archetype. Focusing on 137 cancer-relevant compounds with defined targets across nine CRC cell lines (**Fig. 5c**), we mapped how targeted therapies bias cell-state trajectories toward specific phenotypic endpoints.

We found 54 perturbations had the highest probability of pushing cells towards malignant-related archetypes (archetypes 5, 11, 12, 13, and 15). Archetype 9, a normal cell with an intestinal stem cell (ISC)-like gene expression programme, was the most frequently induced by perturbations (n = 19 perturbations) with a maximal likelihood of inducing transitions (range 0.18 - 0.99) followed by archetypes 13 and 15 which have a similar ISC-like programme but are primary tumour cells (**Fig. 5c**). Conversely, archetypes 6 (EMT programme) and 7 (osteoblast programme), representing normal cells, showed the greatest number of perturbations with minimal transition probability towards them (28 and 31 perturbations, respectively; **Fig. 5c**). Our analysis suggests cells with a high probability to transition towards archetypes with the ISC-like programme following perturbation may exhibit an adaptive stress response, consistent with a ISC phenotype observed in CRC cells resistant to chemotherapy^42^.

We observed both convergent and divergent transition profiles across perturbations. Some drugs with similar mechanisms produced similar state shifts—for example, pan-RAS inhibitors (BI-3406, RMC-6236) and RAF inhibitors (vemurafenib, encorafenib) convergently drove cells toward archetype 14, a pre-malignant cell with a neuroendocrine-like state. In contrast, mechanistically related agents also diverged: dabrafenib preferentially induced transitions toward archetype 15 (primary tumour cell, ISC-like programme), whereas downstream MEK inhibitors favoured archetypes 3 and 4. Conversely, drugs with distinct mechanisms could converge phenotypically; a subset of EGFR/ERBB2 and mTOR inhibitors promoted transitions toward normal states (archetypes 7 and 10), while topoisomerase inhibitors, including irinotecan, consistently biased cells toward tumour archetypes 11–13 (**Fig. 5c**). Quantifying transition probabilities toward defined archetypes thus reveals both mechanistic convergence and divergence in how therapies reshape cancer cell states.

We next analysed cell-state transitions induced by all Tahoe-100M perturbations across three *BRAF*-mutant CRC cell lines, enabling assessment of the context specificity of perturbation-driven state changes. We focused on archetypes 1, 3, and 15, representing normal, pre-malignant, and malignant states, respectively, and together spanning the phenotypic diversity of the Epi-CRC atlas. In aggregate, perturbations preferentially shifted RKO cells along the normal–tumour axis (archetypes 1 and 15; **Fig. 5d**, left), drove COLO 205 cells predominantly toward normal states (archetype 1; **Fig. 5d**, middle), and positioned HT-29 cells along the normal–pre-malignant axis (archetypes 1 and 3; **Fig. 5d**, right). This context dependence was consistently observed across all nine CRC cell lines in the Tahoe-100M dataset (**Supplementary Note 4**). These results reveal cell line–specific responses to perturbation, with the potential to redirect malignant cells toward pre-malignant or normal phenotypic states in a context-dependent manner.

### MAPK inhibition shifts cells towards converging cellular fates

Mapping perturbations onto the Epi-CRC atlas could enable perturbation response modeling to specify therapeutically relevant cell-state transitions. To test this, we prioritised the inflammatory (NF-κB) state (state 33)—given its positive association with stable disease both pre- and on-treatment (**Fig. 3c**)—and the stem, TA-like proliferative state (state 34), which is associated with poorer outcomes on-treatment, as opposing phenotypic targets. We initially focused on *BRAF*-mutant HT-29 cells, which exhibited transitions along the normal and pre-malignant archetype axes (**Fig. 5d**), indicating potential accessibility toward these states. In HT-29 cells, EGFR and KRAS inhibition predominantly shifted cells toward the stem, TA-like proliferative state (state 34), a less favourable phenotype (**Fig. 5e, f**), whereas BRAF inhibition produced variable effects, with dabrafenib inducing a strong shift toward the inflammatory (NF-κB) state (state 33), while encorafenib and vemurafenib elicited modest transitions.

The patterns of cellular transitions induced following dabrafenib treatment were similar in COLO 205 but this was not observed in RKO (**Extended Data Fig. 6a**). RKO exhibited the least sensitivity towards dabrafenib (raw IC_50_ = 4.22 µM) among the three *BRAF*-mutant Tahoe-100M CRC cell lines, potentially reflecting its modest patterns of cellular transitions following treatment with dabrafenib (**Extended Data Fig. 6a, b**). Our findings further support convergent and divergent responses found in the RAF inhibitors and cell line–specific responses to perturbation (**Fig. 5c, Supplementary Note 4**).

We then used our patient-derived organoids as an independent cohort capturing disease relevant cell states. Specifically, we performed scRNA-seq on cells from *BRAF*-mutant (COLO_027) and *KRAS^G12C^*-mutant (COLO_013 and COLIVM_005) CRC organoids following 24 hours of treatment with encorafenib (BRAF inhibitor) and sotorasib (KRAS G12C inhibitor) respectively to validate our reference-guided approach for modelling cellular transitions along identified axes of cell states (**Fig. 5e**). These specific inhibitors were selected on the basis of their clinical use for patients with CRC^52,53^ and we confirmed the selective sensitivity of organoids to their respective targeted inhibitors (**Extended Data Fig. 6c**). We additionally included the hybrid - endoderm-like 2 state (state 21) as an axis to model perturbation responses of MAPK inhibition given the importance of adaptive plasticity towards MAPK inhibitors in both *KRAS*- and *BRAF*-mutant CRC^13,54,55^.

We found that MAPK inhibition across the CRC organoids converged primarily in shifting cells along the axis of the stem, TA-like proliferative (state 34) and the hybrid - endoderm-like 2 states (state 21). Shifts towards the inflammatory (NF-κB) state (cell state 33) were modest across the CRC organoids following MAPK inhibition, suggesting a likely secondary response among the three cell states of interest **(Fig. 5g**). We validated our reference-guided framework orthogonally using a combination of marker gene expression, the proCSC and revCSC signatures^47^, and cell type *k*-NN label transfer from the atlas. Concordant with our predictions, we found marker genes of proliferation (*MKI67, TOP2A*) and MAPK signaling (*DUSP6, ETV4, ET5, SPRY4*)^6^ decreased while marker genes of endoderm development increased (*PROX1, NKD1, SOX4, BMP4*)^11^ following MAPK inhibition in the organoids (**Fig. 5h**).

Cell type *k*-NN label transfer is consistent with our transition analysis as the organoids exhibited a shift to the hybrid - endoderm-like 2 state (state 21) while the proportion of stem, TA-like proliferative state (state 34) cells decreased following perturbation (**Fig. 5i**). Irinotecan-treated Moorman organoids had similar cell state shifts towards the hybrid - endoderm-like 2 state (state 21, **Extended Data Fig. 7a**). The proCSC score was higher in the DMSO samples across the three CRC Sanger organoids (**Extended Data Fig. 7b**), whereas the revCSC score was slightly elevated following MAPK inhibition (**Extended Data Fig. 7c**). Our results support previous findings^11^ and suggest that stress in epithelial cells caused by targeted inhibition and chemotherapy shifts cells towards cell states with developmentally primitive features^11,13^. Together, we demonstrate the possibility of reference-guided modeling of perturbation responses, including convergent mechanisms as a result of MAPK inhibitors. This represents a way forward to generate and test therapeutic hypotheses through integrating both observational and perturbational scRNA-seq data.

## Discussion

We present a continual learning (CL) framework that incrementally expands cell atlases to more accurately capture emergent features of CRC progression as datasets grow. Unlike existing atlas-scale integration solutions^31,37^, which are prone to over-integration^56^, our expansion by regularization approach balances the alignment of healthy states while retaining context-specific variation. Applying this framework, we expanded a published integrated reference atlas with case-control CRC datasets to build a comparative CRC atlas spanning normal, pre-malignant, primary tumour, and metastatic samples, while preserving both healthy and patient-specific disease variation across individuals and studies. This enabled systematic analysis of cell-states across the malignant continuum, is critical for precision medicine approaches based on patient stratification, and is a definitive resource for comparative CRC biology.

Single-cell RNA-seq studies typically classify CRC cell types based on marker genes and transcriptional similarity to healthy colon epithelium^1,3,11^. However, transcription factor (TF) activity has emerged as a powerful approach to delineate malignant subpopulations and their regulatory drivers^3,11,57^. We identified multiple malignant cell states with distinct transcriptional and TF programmes that do not fully resemble non-malignant epithelial lineages, which we term “hybrid” states. Among these, endoderm-like states were enriched in microsatellite-stable (MSS) CRC, whereas microsatellite instability–high (MSI-H) tumours were enriched for an inflammatory (NF-κB) state (cell state 33). Both states exhibited high Shannon entropy and were widely shared across patients, suggesting they represent core features of CRC biology.

Atlas-scale immune–epithelial homing analysis revealed that malignant epithelial states preferentially associate with exhausted T cells, CD16⁺ NK cells or IL10⁺ regulatory T cells. The hybrid endoderm-like 1 state (cell state 20) showed the strongest affinity for exhausted T cells and was positively associated with *KRAS* mutation and progressive disease. CRC organoids recapitulated the hybrid endoderm-like 2 state (cell state 21), linked to *KRAS* mutation, MSS CRC and oncofetal features, supporting their suitability as models of malignant reprogramming. Together, this comparative atlas connects malignant cell states with regulatory programmes, immune context and clinical features, providing a resource to contextualise pre-clinical models and patient outcomes.

Our study establishes a reference-guided framework that directly links a comparative observational disease atlas with an interventional perturbation atlas. By mapping the Tahoe-100M dataset onto our CRC atlas using relative representations, we enable systematic modelling of perturbation-induced cell-state transitions. We anticipate that replacing absolute with relative representations for contextual encoding, as implemented here, will enhance the generalization of generative models for perturbation response prediction^58,59^. Notably, perturbations induced cells across axes of multiple archetypes, underscoring transitions occur along axes of cellular states, the broad accessibility and plasticity of cancer cell states, and revealing substantial opportunities for their rational modulation. We find that perturbations drive coordinated shifts along continuous phenotypic axes rather than toward isolated discrete states, in a manner that is context dependent. Using our reference-guided analysis, we identified a converging mechanism of response along the axis of the stem, TA-like proliferative (state 34) and the hybrid - endoderm-like 2 states (state 21) in CRC organoids following MAPK inhibition.

The variable penetrance and expressivity of post-perturbation cell transitions highlight the richness and complexity of the underlying gene regulatory networks, pointing to fertile ground for discovery. We observed divergent and convergent drug-induced state transitions, including for drugs with the same nominal target, pointing to the utility of perturbation-based approaches for differentiation of drug activity, including mechanism of action, potency, selectivity and safety. How much transcriptional or compositional cell state shifts translate into meaningful clinical endpoints requires further interrogation. Future integration of richer clinical annotation with tissue single-cell multi-omics and spatial profiling, alongside reference single-cell atlases of genetically and pharmacologically perturbed organoids that faithfully capture tumour cell states, will sharpen predictive cell-state modelling and accelerate the development of rational cell-state–directed therapies.

In conclusion, by directly linking a comparative observational atlas with a perturbational atlas, our framework shifts atlas-based single-cell analysis from descriptive state annotation toward predictive and interventional modeling. We anticipate that such reference-guided approaches will enable causal effect modeling^12,60^, rational manipulation of malignant cell states across cancers, and accelerate the development of cell state-targeted therapies.

## Methods

### Continual Learning for extending single-cell atlases to case-control data

Continual Learning (CL) addresses the challenge of training neural networks on streams of data where distribution shifts across the sequence violate the Independent and Identically Distributed (I.I.D) assumption. Differently from Transfer Learning (TL), where network components are frozen for information transfer, CL maintains full model trainability while employing mechanisms (e.g., regularization^38^, replay^61^, dynamic network expansion^62^ to balance stability and plasticity (adaptability). These strategies mitigate Catastrophic Forgetting (CF) while preserving the capacity of the model to adapt to emerging tasks and maintain competitive performance across both past and novel tasks.

A common practice in cell atlasing efforts is to *extend* integrated healthy reference atlases to include disease samples via Transfer Learning, also known as architecture surgery^37^. Inspired by increasing interest in continually extending the integrated reference atlases, we hypothesized that where the intention is to introduce a *major update* to an existing integrated cell atlas by adding heterogeneous disease datasets, Continual Learning improves existing atlas extension practices.

Motivated by the findings by Dann *et al*.^63^, who found that disease cell states are best preserved by TL when the query contains case and control samples, we introduce a regularization for continual expansion of reference cell atlases with case-control query data. Our regularizer combines Experience Replay and a modification of Elastic Weight Consolidation (EWC) tailored to case-control data to improve preservation of novel disease states, prevent forgetting and improve model adaptability.

### Elastic Weight Consolidation regularization for case-control data

Let *θ* denote the weights in an encoder-decoder-based neural network (i.e., Variational Autoencoders) trained on input data *X*. Under Gaussian initialization, *θ* follows a normal distribution. Then the density is given by

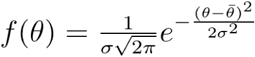

The Elastic Weight Consolidation (EWC) penalty^38^ is a common technique to prevent catastrophic forgetting in CL. Let *θ^OLD^* A common practice in cell and *θ^NEW^* be respectively the parameters of the old and new model in a continual training. Let *F* denote the Fisher Information for the parameters in *θ*. EWC is defined as

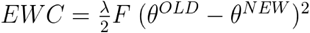

This suggests that EWC regularization in a continual learning setup is equivalent to defining a Gaussian prior for the weights in the new model, centered on parameters in the old model with a precision defined by *F*^−1^ , the inverse Fisher Information. That is,

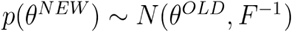

In continual comparative mapping, we let *F* = F*_θref_* O F*_θ_*_quary ctrl_, be the Hadamard product of *F*_θ*ref*_, the Fisher Information with respect to healthy cells in the reference (atlas) model, and *F*_θquery crtl_, the Fisher Information computed with respect to a proportion of cells sampled from query control cells. That is, in comparative mapping we place a prior on the weights of the query model as follows:

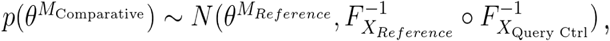

Where *X_Reference_* is a subset, usually 20%, of the cells from the datasets in the reference cell atlas (also called the Replay buffer) and *X*_Query Ctrl_ is a subset (5%-10%) of the control cells in the query data.

### Experience Replay for improved adaptability

Experience Replay (ER) involves revisiting samples from previously observed data during training. To enable this, a Replay buffer stores a selected subset of data, typically 10%-30% of the total, for rehearsal. By replaying past knowledge along with new data, the approach prevents feature drift and improves model adaptability. In this work, the buffer stores a subset of cells from the reference atlas datasets, selected either randomly or according to a specific criterion. While the main results in the manuscript rely on random selection, we benchmark random selection and a guided selection strategy based on predictive uncertainty (see “*Bregman Information for optimal selection of Experience Replay”*).

### A regularizer for continual atlas expansion with case-control data

We compute the total loss by combining the modified EWC and Replay as

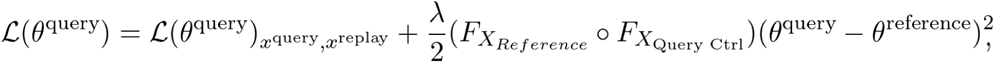

where 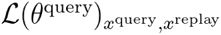 is the loss (ELBO) from the base model computed on both query and the Replay buffer. Here, the reference model was trained using scANVI^64^. The loss here, therefore, is the loss of the scANVI model.

Concurrent to our work, Daxberger *et al*.^65^ proposed a similar, yet different, regularization and demonstrated improved Continual Learning. Note that unlike Daxberger^65^ who computes EWC and Replay from disjoint subsets of the past data, we compute *F_XReference_* on the Replay buffer. Therefore, the selection of cells in the Replay buffer can be important. In the next section we introduce alternative strategies for selection of cells (experiences) in the Replay buffer that are based on a generalised variance of the latent space that we compute by estimating the Bregman Information (BI)^66^. We compare random selection and our alternative selection strategy, which we refer to as BI hereafter, for different buffer sizes in **Supplementary Note 3, Supplementary Fig. 2b-d**.

### Bregman Information for optimal selection of Experience Replay

Model uncertainty can be disentangled into aleatoric (inherent in the data and irreducible) and epistemic (reducible by collecting more data)^67,68^. To obtain a sample-wise estimate of epistemic uncertainty, Gruber and Buettner^69^ introduced a bias-variance decomposition of loss functions that are proper scoring rules. They demonstrated that the variance component of the cross-entropy loss has the closed form of a Bregman Information (BI) generated by the Log-Sum-Exp (LSE) function acting on the model’s logits and can be directly related to the epistemic uncertainty. Serra *et al*.^70^, further demonstrated that using a BI-based estimate of a model’s epistemic uncertainty to populate the Replay buffer in memory-based Continual Learning alleviates Catastrophic Forgetting (CF). Following Serra 2025, we estimate BI by approximating the required expectations with a finite ensemble obtained via test-time augmentation (TTA)^71^ and use the resulting estimates of epistemic uncertainty for memory management and reducing CF.

Specifically, for a given cell *x*, we employ the TTA to generate *J* perturbations of the cell. We encode the latent representations 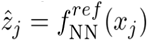 of the perturbed cell *x_j_* using the encoder of the reference model, 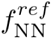, and estimate the BI as follows:

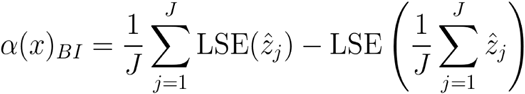

where 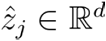 and 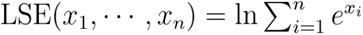 are the latent encodings and the Log-Sum-Exp function respectively. The perturbed latent representations are generated by randomly masking 50% of the genes in the cell^72,73^, then using the encoder to get 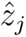.

Large values of BI imply that the generalised variance of the latent representations of a cell across perturbations is large, thus the model is not confident about the considered cell (high epistemic uncertainty). In **Supplementary Note 3,** we study the effect of three Replay buffer selection strategies on the distance of healthy replayed cells from the reference atlas: bottom-k (most representative cells with low BI), top-k (most uncertain, high BI), and mixed sampling (combining cells from both bottom-k and top-k) (**Supplementary Note 3, Supplementary Fig. 2b**). We also consider the distance of a simulated novel biological state that is only present in the case (disease) of the case-control query data after CL training (**Supplementary Note 3, Supplementary Fig. 2d**). The regularization and Replay buffer selection schemes developed here are implemented for scANVI models scvi-tools version 0.16.1 (**Code Availability**).

### Quantification of cell state transitions upon perturbation using Relative Representations and Energy Distance

Relative Representations^48^ were originally proposed for aligning the latent representations of two pre-trained models. A set of anchor points 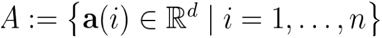 are randomly selected from the observations. For each observation their encoding is computed. Then, new coordinates are defined by computing the cosine similarity between the absolute representations of each observation in the dataset and those of the anchor points:

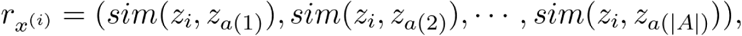

which define the Relative Representations **r**_x(i)_. Here *sim*(.,.) denotes the cosine similarity between two representations.

In this work, we select the anchors from archetype or cell state neighbourhoods in the CRC atlas, which represent a target cell state. We then compute the relative representations for each cell from the pre-clinical model samples, organoid or cell line, and quantify state transition upon a perturbation by comparing the relative representations pre- and post-treament via Energy Distance.

Let *X* ∼ *P* and *Y* ∼ *Q* be random variables and let || . || denote the euclidean distance. The Energy Distance ε(*P, Q*) is defined as:

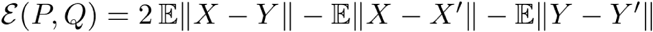

Let 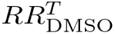 and 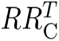 denote the Relative Representations of DMSO/unperturbed and chemically perturbed cells in condition *C* to a target cell state *T* respectively.

To quantify the shift induced by an exogenous perturbation such as a small-molecule compound towards a desired cell state, we compute 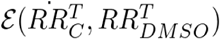. We then obtain the probability of transitioning to a state *T* by normalizing the E-distances over multiple states *T* = *T*_1,_ *T*_2, … ,_ *T_m_* by applying a temperatured softmax as follows:

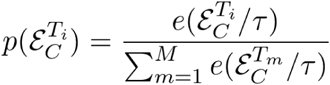

While not covered here, a p-value can be assigned to state transitions using the permutation-based framework of Peidli *et al*.^74^ for assigning statistical significance to Energy Distance between the control and perturbed state.

### General data preprocessing

#### Public Single-cell RNAseq data and Tahoe-100M

We used the count data supplied by the original studies. Please refer to the original publications for details of preprocessing. Each dataset was subsetted to the set of genes in the reference model from Oliver *et al*. and was padded with zeros for non-overlapping genes.

The Tahoe-100M dataset^28^ was downloaded as per author instructions. Raw counts for all available CRC cell lines (n=9) were subsetted to the set of genes in the reference model from Oliver *et al*.^29^ and were padded with zeros for non-overlapping genes.

### PBMC simulations

We simulated single cell datasets to represent two major biological events associated with disease occurrence that can be inferred from case-control scRNAseq data^75^ cell type composition shifts and transcriptional shifts.

#### Novel disease-specific state simulation

To simulate cell type composition shifts, we followed the workflow proposed by Dann *et al.*^63^ for introducing out-of-reference (OOR) cell types into single-cell datasets. These simulations aim to model scenarios where specific cell type clusters are removed from the healthy reference datasets, resulting in compositional changes. We used the preprocessed and harmonized healthy PBMC 10X Genomics scRNA-seq data curated by Dann *et al.*^63^.

To construct the reference atlas, control, and pseudo-disease datasets, we followed the strategy outlined by Dann *et al*.^63^. Donors were selected from the one query study and split randomly into disease and control subsets. We designated the remaining 12 studies, comprising 1,219 donors, as the atlas dataset. Composition shifts were then introduced by dividing Naive B Cells and T Cells into OOR and in-reference groups. To achieve this, we first log-normalized the gene expression profiles of the selected cell type in the disease and control datasets. Using principal component analysis (PCA), cells were then split into OOR and in-reference groups based on their weights along the first principal component. To generalize this distinction across datasets, we trained a *k*-nearest neighbour (*k* = 10) classifier on the PCA-based splits in the query dataset and used it to assign atlas cells to the corresponding groups. This method allowed us to simulate variable OOR states. Importantly, by using PCA, we ensured that the first principal component captured biologically meaningful variation within the query dataset, avoiding issues related to batch effects.

Naive B Cells were chosen for their homogeneous transcriptional profiles, making them an ideal candidate for initial experiments involving straightforward compositional changes. Removing them allowed us to create a clear and interpretable OOR state. On the other hand, T cells offered a relatively more challenging test case due to their inherent heterogeneity. As one of the largest and most diverse cell populations profiled, T cells encompass multiple distinct states within a single cell type. This complexity made them an excellent candidate for evaluating how well our methods perform under biologically realistic and diverse conditions.

The simulated composition-shifted datasets included a reference atlas representing the healthy dataset without OOR states on which the reference scANVI model was trained, a “control” dataset mimicking healthy profiles with excluded OOR states, and a pseudo-disease dataset enriched with OOR cell states to simulate disease conditions.

#### Simulation of shift in the IFN signature in Monocytes

To simulate transcriptional shifts in monocytes reflecting an interferon (IFN) response, we used the same preprocessed and harmonized healthy PBMC 10X Genomics scRNA-seq data curated by Dann *et al*., that we used for composition shift simulation. Using the preprocessed dataset, the following steps were taken to simulate IFN-stimulated monocytes: We built a reference from the preprocessed PBMC dataset, 10 donor IDs were randomly selected and removed. The remaining dataset, after excluding these 10 donors, was designated as the atlas, used for training the reference scANVI model. The 10 removed donor IDs were split into two groups. 5 donor IDs were assigned to the control group and 5 donor IDs were assigned to the disease group.

To simulate the disease group, we utilized the scDesign3^76^ pipeline, with the dataset corresponding to the five disease donors serving as input. First, marginal models were constructed using the scDesign3 framework, where the mean (*μ*) parameter of gene expression was modeled as a function of the interferon (IFN) score. This allowed for the incorporation of the IFN score as a biologically relevant covariate influencing gene expres- sion patterns. Next, a Gaussian copula was fitted to capture the correlations between genes, thereby preserving the dependencies and interactions observed in real biological systems.

To simulate a heightened transcriptional response characteristic of disease conditions, the coefficients of the IFN score in the marginal models were doubled. This adjustment reflected an elevated influence of the IFN score on gene expression, mirroring the transcriptional shifts associated with disease states. Synthetic gene expression data for the disease group were then generated by sampling from the Gaussian copula, and the sampled values were transformed using the adjusted marginal models to ensure biologically realistic characteristics. The final simulated dataset included a reference containing the remaining donors after removing the 10 donor IDs, a “control” dataset representing the original gene expression data for the 5 control donors, and a disease dataset containing the simulated transcriptional profiles for the 5 disease donors generated by scDesign3. All scVI/scANVI models used for construction of the reference, de-novo integration and architecture surgery were trained using scvi-tools (version 0.16.1).

### Replay buffer management experiments

Using the simulated data described in “*Simulation of shift in the IFN signature in Monocytes*”, we benchmarked the importance of Replay buffer characteristics to the plasticity (adaptability) of the model. We considered Replay buffer sizes of 10%, 20% and 40% , where cells to replay were selected randomly or guided by the BI metrics through the “top-k”, “bottom-k” and “step” schemes described earlier. We repeated each experiment 10 times. The experiments were subject to an 8-hour wall-time limit.

We used two metrics to evaluate the performance of the model under each experiment setup. We use L2-normalised squared Wasserstein distance 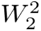 between replayed cells and reference as a measure of distortion to the original cell representations in the reference post CL. We sample 2000 cells from the replay and reference cells each and compute the squared Wasserstein distance. We repeat this 50 times and compute the mean of 50 replicates, and apply L2 normalisation to make different experiments comparable. The 2-wasserstein distance squared is defined between two distributions z and z’ as:

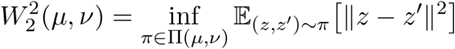

We additionally assess the distance of the novel disease-specific state (increased IFN response signature) to the query control cells as a proxy for preservation of novel states. Specifically this distance is computed as:

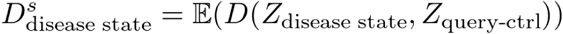

where D(.,.) is the euclidean distance, Z_state_ and Z_query-ctrl_ are the cell representations for simulated disease state and control cells in the query. Here, distances are averaged to compute the final score.

### Comparative all-lineage, Epithelial-lineage and NK-T lineage integrations

For continual expansion of lineage-specific atlases provided by Oliver *et al*.^29^, we are restricted to the architecture of the reference model. Please see Oliver *et al*.^29^, for details on model architecture namely, number of layers, number of latent dimensions, choice of likelihood for gene count reconstruction, batch and additional covariates. The parameters for the final integration were determined based on benchmarks from the simulated PBMC dataset. The following parameters were used:

#### Comparative all-lineage

This model was trained using a Replay buffer generated by random selection of 20% of the cells in the all-lineage (healthy Pan-GI) integration reference model. EWC importance was set to 100. The model was trained for 100 epochs. The model was conditioned on the study due to heterogeneity in cancer cell states across the datasets.

#### Comparative Epi-CRC

This model was trained using a Replay buffer generated by random selection of 20% of the cells in the Epithelial (Large Intestine Adult-Pediatric) integration reference model. EWC importance was set to 100. The model was trained for 200 epochs. The model was conditioned on the study due to heterogeneity in cancer cell states across the datasets.

#### Comparative Immune-CRC

This model was trained using a Replay buffer generated by random selection of 20% of the cells in the T and NK cell integration reference model. EWC importance was set to 0.5. The model was trained for 200 epochs. The model was conditioned on the DonorID as we aimed for capturing common states across datasets.

### Benchmarking Epi-CRC integration

We calculated distance metrics to evaluate the cellular shifts after integration. The “shift cancer” metric is the ratio of the distance between tumour (case) sample to the normal (control) sample and to the distance between normal sample in the query and healthy samples in the reference. “Shift control” is the distance between query-control and reference samples. In addition, we also calculated ordering and divergence scores to evaluate how cells are ordered in known trajectories and how much they deviate from healthy trajectories as detailed in Nazaret, Fan, and Lavallée *et al*.^17^. We compute these metrics on the comparative embedding (i.e. latent space of the integrated datasets). We use these metrics in replacement of the conventional scIB metrics^77^ as scIB metrics are primarily designed and tested to evaluate integration of homogenous healthy cells.

#### Shift-control

This metric measures how the cross-distances of normal cells in the query and reference compares to the average of within reference and within query distances. Distances are computed on the latent space and averaged. A small score indicates good mixing between normal cells in the query and reference.

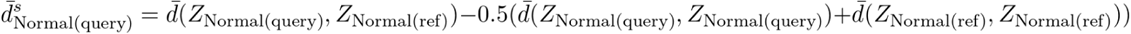

#### Shift-cancer

This metric measures how large the cross-distances between tumour and normal cells in the query are relative to the distance between Normal cells in the query and the reference. It suggests how well the transcriptional shifts due to the disease (cancer) are captured in the comparative embedding.

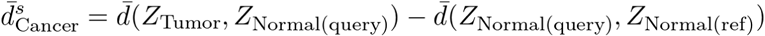

Distances are averaged to compute the final score. A large value indicates disease shifts are captured well in the comparative embedding.

#### Shannon Diversity Index

This metric measures the mixing of cells from S different groups, donor in our case, in a dataset. We compute SDI per leiden cluster or cell type. A large value indicates cells from many patients are represented in a given group, whereas a low value indicates the group contains cells of few or one individual.

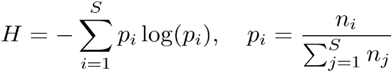

#### Divergence score

This metric measures the difference between mean distances of two non-neighbour groups and neighbour groups (e.g. cell states or progression stages) having different origins in a trajectory. This metric is a measure of divergence from the trajectory. We consider the malignancy progression trajectory to compute this score. Distances are computed by randomly sampling cells from CRC stages (normal, primary, metastatic). A large value suggests that the divergent branches in a pre-defined trajectory are preserved.

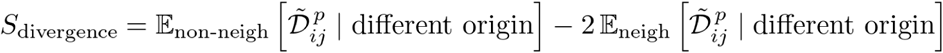

#### Ordering score

This metric is the ratio of distances between cells from non-neighbour and neighbour stages that share the same origin in the trajectory. A large value indicates that the order of cells along a pre-defined trajectory is preserved.

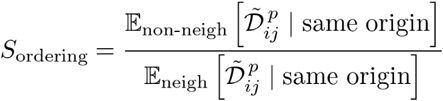

### Cell type and cell state annotation

#### Cell type and cell state annotation of the comparative Epi-CRC Atlas

To annotate disease cell types and states in the comparative Epi-CRC atlas, we first performed leiden clustering on the comparative embedding. We used a clustering resolution of 0.1 for pre-malignant cells. For malignant cell types and states, we performed clustering using resolutions ranging from 0.5 to 2.0 with increases in the resolution parameter of 0.1 and calculated the Adjusted Rand Index (ARI) for each clustering resolution. We selected a clustering resolution of 1.0 for the final number of clusters as the ARI values were consistently > 0.8 on a heatmap, generating 49 clusters for annotation.

We used a combination of cell type markers from Elmentaite *et al*.^34^ and Pelka *et al*.^1^ initially to label cell types in the Epi-CRC atlas. For the remaining clusters that did not fit gene expression profiles of the marker genes, we used transcription factor (TF) activity scoring with Decoupler (version 1.5.0) to obtain a TF activity score per cluster. We manually consolidated each identified TF for the final cell type and cell state annotations in the comparative Epi-CRC atlas. For hybrid cell states that are challenging to annotate, we used ChatGPT-4 to cross-check TFs. We verified all final annotations and their interpretations. To supplement the TF analysis, differential gene expression (DGE) analysis was performed using a Wilcoxon test (sc.tl.rank_genes_groups, method=’wilcoxon’).

#### Cell type annotation of the comparative Immune-CRC Atlas

We used a weighted *k*-NN prediction approach to annotate the T and NK cells in the comparative Immune-CRC atlas. We used the T and NK cells from Oliver *et al*.^29^ as the reference dataset to fit a *k*-NN search tree using Pynndescent (version 0.5.1) and the remaining cells from the assembled cancer datasets were designated as the query.

We first created a *k*-NN index using the reference data and the *NNDescent* function. We used the embeddings of the query cells to obtain the distances of nearest cells from the reference to each query cell using the default number of neighbours (*k* = 10). We converted these distances into weights using a Gaussian Kernel. Cells from the query with a smaller distance to the reference have a larger weight given neighbour approximation towards the cells in the reference. Finally, we assign the final label to each cell in the query by assigning the cell type label from the reference with the highest sum of weights. To confirm the final cell type labels, we consolidated marker genes from CellTypist (https://www.celltypist.org/encyclopedia/Immune/v2).

### Immune homing analysis

We used the cell homing analysis toolkit cell2home (To, Pett, Dufva *et al.* in preparation) to predict homing interactions between epithelial cell states and T/NK immune cell subsets.

The epithelial cell object was used to pseudobulk cell states and calculate a reference of chemoattractant ligand expression levels across the epithelial states. This reference was then used to compute homing affinity scores for each immune cell in the T/NK object towards the epithelial states, by aggregating chemoattractant receptor-ligand interaction scores into homing affinity scores. For visualisation as a heatmap, homing affinity scores were averaged for each immune cell type and scaled across the immune cells. Ligand-receptor pairs with strongest contribution to the homing affinity score between cell types of interest were ranked based on high expression of both the receptor in the migrating cells and the ligand in the attracting cells.

### CRC organoid scRNA-seq experiments and data processing

#### CRC organoid culture and drugging

We cultured CRC organoids from the Wellcome Sanger Institute’s Cellular Generation and Phenotyping core facility. We cultured the CRC organoids using the media components found in **Supplementary Table 4**. We used a mixture of 80% Basement Membrane Extract (BME) and 20% organoid media for culturing the CRC organoids^78,79^. We passaged the organoids an average of once weekly once they reached 80-90% confluency.

For the drugging experiments for scRNA-seq profiling, we plated the organoids at a density of 1,000,000 cells per well in a 6-well plate (Corning). Following 72 hours of incubation to allow organoid reformation, we treated the organoids with the vehicle control (0.1% DMSO) and small molecule inhibitors as described in **Supplementary Table 4**. After 24 hours of incubation with the vehicle control and drugs, we dissociated the organoids chemically by suspending them in TrypLE and incubation in a 37°C water bath for 10-15 minutes. We also mechanically dissociated the organoids with the P1000 and these processes were repeated until we obtained single-cell suspensions. We used the protocol from 10x Genomics (Chromium Single Cell 3′ Gene Expression v.3.1 Chemistry Dual Index) to perform cellular barcoding using the 10x Chromium. The Wellcome Sanger Institute’s DNA Pipelines core facility performed library preparation followed by paired-end sequencing using the Illumina NovaSeq 6000 platform with 150 bp paired-end reads.

#### CRC organoid data processing and filtering

The Wellcome Sanger Institute’s New Sequencing Pipeline group aligned and mapped sequencing reads to the human reference genome (GRCh38) using CellRanger (version 7.0.0) to generate an unfiltered count matrix. The Wellcome Sanger Institute’s Cellular Genomics IT team performed sample demultiplexing using Souporcell^80^ and genotype data for each organoid.

We performed initial quality control on the unfiltered count matrix by first calculating the inflection point of ranked barcode counts on a cumulative sum curve^81^ and filtering cells with less than 200 genes expressed. We identified outlier cells that are defined as 5 median absolute deviations (MADs)^82^ of *log1p_total_counts*, *log1p_n_genes_by_counts*, and *pct_counts_in_top_20_genes* calculated using the calculate_qc_metrics function in Scanpy. We performed normalization and log raw transformation of raw gene expression counts using Scanpy. We selected 2,000 highly variable genes (HVGs) and designated the organoid line as the batch. We manually included a list of genes as HVGs that are related to colonic cell types and colon function (**Supplementary Table 5**).

We scaled each gene to unit variance (*sc.pp.scale* function) before dimensionality reduction with PCA, retaining PCs that explained 70% of variance in the data before performing leiden clustering to identify poor quality cells. We finally removed outlier cells, doublets designated by Souporcell, and poor quality cells that cluster together before filtering genes expressed in less than 3 of the remaining cells.

### Cell type annotation of Tahoe-100M cell lines and CRC organoids

We first generated relative representation embeddings of the comparative Epi-CRC atlas, Tahoe-100M cell lines, and CRC organoids with 2,000 anchors. The anchors were selected at random from the Epi-CRC atlas. The number of anchors was set to 2000 to capture the heterogenous nature of the reference as much as possible. We then concatenated all three 2,000 dimensional embeddings before label transfer by weighted *k*-NN prediction as described in the “*Cell type annotation of the* comparative *Immune-CRC Atlas”* section above for cell type label transfer using the Epi-CRC atlas as the reference dataset. To quantify uncertainty for the cell type labels, we subtract the sum of weights from 1 to obtain an uncertainty score per cell. Cells with an uncertainty score greater than 0.5 are considered cells with labels with a high degree of uncertainty.

### CRC organoid viability assays

We modified the plating method from Francies *et al*.^79^. We first covered the wells of a 96-well plate (Corning) using a 50 µL mixture of 50% BME and 50% organoid media before 30 minutes of incubation in a 37°C incubator to allow the polymerization of BME. We added ROCK inhibitor to the plating media at a dilution factor of 1:1000. We plated dissociated organoids in 100 µL of organoid media at a density of 2,500 cells per well followed by 24 hours of incubation after plating prior to compound screening.

We treated the organoids using a format of a 9-point dose response curve with 2-fold dilutions for the drug treatments. The maximum concentration was 2 µM, 5 µM, and 10 µM for sotorasib, dabrafenib, and encorafenib respectively. We measured cell viability using fluorescence from CellTitre-Glo (50 µL per well) following 72 hours of incubation in the drug.

### Drug sensitivity analysis

We normalized the raw viability using the negative and positive controls for each tested drug per organoid (averaged across three technical replicates). The measured average fluorescence of the wells with media and cells represented the negative controls. The positive controls were the measured average fluorescence of two wells with media only at the beginning and end of each row. We calculated the viability using the following equation:

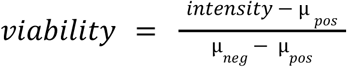

We used technical replicates to fit individual curves and assumed a sigmoidal curve to fit dose-response curves. We used non-linear regression (broom.mixed R package) for curve fitting. We scaled the drug concentrations to compare between different drugs (maximum concentration at 9 as a 9-point dose-response curve was used) as follows:

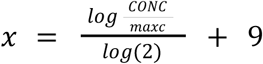

We calculated the mean IC_50_ across three biological replicates in each organoid and drug treatment. We visualized the mean raw IC_50_ on the log scale. We obtained drug sensitivity data of the *BRAF*-mutant CRC cell lines from the Genomics of Drug Sensitivity in Cancer 2 (GDSC2) database via the Cell Model Passports website: https://cellmodelpassports.sanger.ac.uk/downloads

### Archetype analysis

We computed 10 diffusion components for normal, polyp and tumour cells individually using scanpy version 1.11.1. We then used the PCHA python package (version 0.1.3) to compute 10, 5 and 5 extreme phenotypic states in normal, pre-malignant and tumour cells respectively.

## Data Availability

The full list of external datasets is available in **Supplementary Table 1**. Raw counts were obtained from GSE166555 (Uhlitz 2021), GSE108989 (Zhang 2018), GSE201348 (Becker 2022), SCP2079 (Tian 2023), SCP1162 (Pelka 2021), GSE178318 (Che 2021), GSE200997 (Khaliq 2022), https://lambrechtslab.sites.vib.be/en/pan-cancer-blueprint-tumour-microenvironment-0 (Qian 2020), https://www.synapse.org/Synapse:syn26844071/files/ (Joanito 2022), https://cellxgene.cziscience.com/collections/a48f5033-3438-4550-8574-cdff3263fdfd (Chen 2021), and https://github.com/dpeerlab/progressive-plasticity-crc-metastasis (Moorman 2024). The CRC organoids data generated for this study are available by request and will be uploaded to public repositories prior to publication.

## Code Availability

Source code for the comparative atlas construction via CL and relative representations is available at https://github.com/theislab/comparative_atlas. The scANVI models of the all-lineage, epithelial lineage and NK-T cell lineage integrations presented here, and the notebooks to reproduce the figures from this manuscript can be accessed from HuggingFace and the GitHub repository.

## Acknowledgements

We would like to thank Dr Thomas Möllenhoff, Dr Mariela Cortés-López, and Dr Luke Zappia for their constructive feedback on initial drafts of the manuscript. We thank Dr Jack Finlay for initial versions of cell type annotations. We thank Steven Leonard from the Wellcome Sanger Institute’s New Sequencing Pipeline group for performing the initial processing of the raw sequencing reads from the CRC organoid 10x Genomics experiment. We would also like to thank Dr Pavel Mazin from the Cellular Genomics IT team for running Souporcell for sample demultiplexing of the CRC organoids. We thank Dr Howard Lightfoot for advice on dose-response curve fitting.

This work was funded by the European Union (ERC, DeepCell - 101054957) and supported by the German Federal Ministry of Research, Technology and Space (BMFTR) under grant no. 01IS18053A. This work was also funded by the Wellcome Trust Grant 206194 (M.J.G.) and Wellcome Sanger Institute Quinquennial Review 2021–2026, Wellcome Core 220540/Z/20/A (M.J.G., T.S.T.). S.H. was partially supported by a German Academic Exchange (DAAD) scholarship. O.D. was supported by an EMBO Postdoctoral Fellowship [ALTF 420-2023] and by UK Research and Innovation (UKRI) through the Horizon Europe funding guarantee for a Marie Skłodowska-Curie Actions Postdoctoral Fellowship [EP/Z533804/1].

The authors have applied a CC BY public copyright license to any author accepted manuscript version arising from this submission.

## Author contributions

S.H. and T.S.T. conceptualized and designed the project with the help of F.J.T. and M.J.G.. S.H. and T.S.T curated and harmonized the datasets and performed formal data analysis. O.D. performed immune homing analysis. K.T., J.P.P, O.D., and S.A.T. provided early access to the cell2home immune homing analysis toolkit. A.O., R.E. and S.A.T. provided early access to the healthy gut cell atlas. S.H. and R.J. performed benchmarking on the continual learning approach. G.R.P. processed the Tahoe-100M dataset. T.S.T performed the CRC organoids experiments with advice from G.P. and M.J.G.. F.J.T. and M.J.G. supervised and provided funding for the project. S.H., T.S.T., F.J.T., and M.J.G. wrote the manuscript. All authors provided assistance with interpretation of the results and provided feedback on drafts of the manuscript.

## Competing interests

R.E is a co-founder and holds equity in Ensocell Therapeutics. In the past three years, G.P. has been a consultant for Mosaic Therapeutics. S.A.T. is a scientific advisory board member of Bioptimus, ForeSite Labs, Xaira Therapeutics, Board observer and equity holder of TransitionBio, a co-founder, consultant and Board Director of Ensocell Therapeutics, a non-executive director of 10x Genomics and a part-time employee of GlaxoSmithKline. F.J.T. consults for Immunai, CytoReason, BioTuring and Phylo Inc., GenBio, and Valinor Industries, and has ownership interest in RN.AI Therapeutics, Dermagnostix, and Cellarity. AstraZeneca, GlaxoSmithKline, and Astex Pharmaceuticals have awarded M.J.G. research grants. M.J.G. is a consultant for Bristol Myers Squibb. M.J.G. is a board director for and equity holder in Mosaic Therapeutics. The remaining authors declare no competing interests.

## Supplementary Note 1: Incremental atlas expansion using a trained cVAE

Incremental atlas expansion involves iterative updates to the cVAE model that was used to construct a reference atlas to integrate additional query datasets from new experiments or studies. Comparative atlas expansion preserves and contrasts the cellular states in the reference and new dataset being integrated, balancing the alignment of shared variation with retention of context-specific states, which is best achieved using case-control query data.

Motivated by these concepts we designed a continual-learning based regularizer for cVAE-based integration models (**Supplementary Fig. 1**) that improves adaptability of the model to new variations in the (case-control) query by mitigating catastrophic forgetting of the past knowledge as it is being trained on new data, and aligning the healthy control cells in the query and reference.

In each incremental data integration step, a subset of cells used to construct the existing reference model are selected at random or via scoring the variance of the model for each individual cell in the reference using the Bregman Information metric (step 1). We call this subset, the Replay buffer, as these cells will be rehearsed at training time along with case-control query data to help the model retain the old variations learned from the previous data. We then compute importance weights (based on Fisher Information - see **Methods**) for each weight parameter in the cVAE model using healthy cells from the reference and control cells from the query (step 2) and penalize the model such that the important weight parameters are protected from hazardous updates that would be detrimental to variation in the reference. The penalization to model weights is computed by EWC metric (**Methods**), which we modified to accommodate the case-control data. Together, the Reply buffer and EWC improve model adaptability and enable continual expansion of the reference atlas.

Technical effects such as batch and study are controlled during continual expansion by the conditioning design of the cVAE models, as in architecture surgery. However, unlike architecture surgery the weights are unfrozen in our incremental update with regularization setting. While no particular order for adding the data is assumed, it is worth noting that the model can become more prone to catastrophic forgetting as the number of incremental updates increases, especially if the regularizer parameters are not sufficiently optimal.

**Supplementary Fig. 1:**
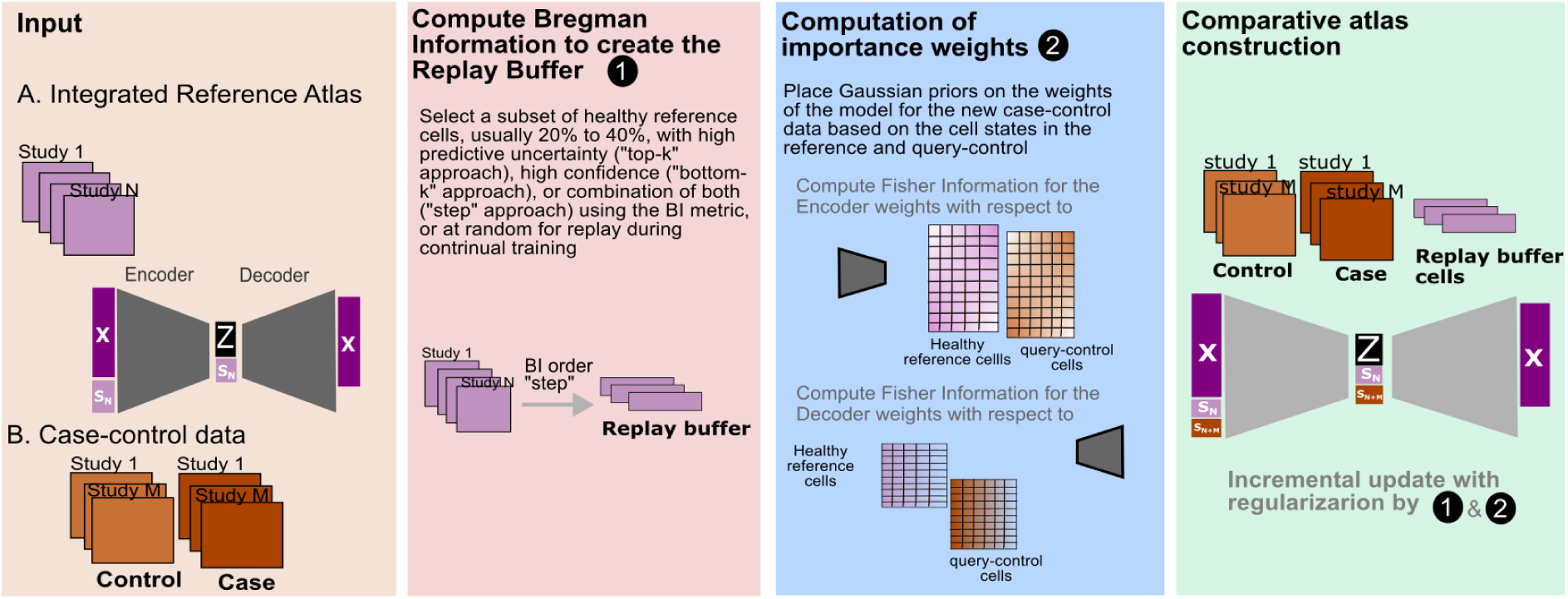
Overview of comparative atlas expansion with case-control data.

## Supplementary Note 2: Interpretation of Epi-CRC benchmark results

The reference from Oliver *et al.*^29^ is a scANVI model. We mapped the ten curated CRC datasets to the reference by architecture surgery using *donor ID* as the conditional covariate for data integration (*scANVI:donor)*. We mapped the CRC data in a separate model using *study* as the conditional covariate (*scANVI)*. These two models serve as our baselines. We applied our CL approach setting the importance of the EWC to 100 and replaying 20% and 60% of the reference cells (*Replay 0.2,EWC100* and *Replay 0.6,EWC100 respectively*). We additionally trained a model without regularization (*EWC0, Replay0*). All models trained by CL were conditioned on *study*. Sample-level embeddings are generated through mean aggregation of cell-level embeddings. Distances were computed on the cell- and sample-level embeddings obtained from this model after architecture surgery or CL.

We found that tumour samples integrated with our CL approach with regularization parameters (*Replay 0.2, EWC 100*) exhibited the largest distance from the control samples and lowest distances between reference and query control samples when compared to integration with architecture surgery (**Fig. 2g**). To ensure that our CL approach re-capitulates disease progression from normal to metastatic states, we used samples from Moorman *et al*.^11^, given the availability of baseline characterization of the transition from normal to metastatic states. Fine-tuning –no replay, no regularization–, that is (*Replay0, EWC0*), showed the highest ordering and divergence scores at the sample level, while CL and *scANVI:donor* performed comparably (**Extended Data Fig. 1b**).

At the cellular level across all cells, *scANVI:donor* showed the largest divergence score while our CL approach maintained a balance in both ordering and divergence scores suggesting that CL training has improved model adaptability (**Extended Data Fig. 1c**). When comparing both methods for integration using only cells from Moorman *et al*.^11^, we found embeddings from our CL approach had both higher ordering and divergence scores compared to the two scANVI baselines (**Fig. 2i**).

## Supplementary Note 3: PBMC simulation benchmarks on Replay buffer management

Our experiments on the importance of the regularization strength of EWC and Replay in simulated data characterised by disease-specific cell type compositional and transcriptional shifts suggests that a model in which reconstruction of the replayed cells is heavily penalized is harmful to the adaptability of the model (**Supplementary Fig. 2a**). Our simulations also suggest that CL outperforms de-novo integration and integration by Transfer Learning (i.e. architecture surgery) when there is a balance between replay and EWC regularization strength and generally for moderate regularization by replay. We, therefore, did not enforce any important weight to the Replay part of the regularizer in our final formulation of the loss function. That is, reconstruction of replayed cells is equally weighted as the reconstruction of cells from case-control query.

We additionally studied the characteristics of the Replay buffer required for a model with high adaptability capabilities. We compared the BI-informed buffer management strategies proposed in this work, that is selection of the buffer through the “step”, “top-k” and “bottom-k” schemes, with random sampling from the reference as our baseline. We ran the integration of simulated data described in “*Simulation of shift in the IFN signature in Monocytes”* for buffers of size 10%, 20% and 40% of the reference cell through each BI scheme. Each experiment was repeated 10 times using EWC importance of 0.5 (**Supplementary Fig. 2b-d**).

**Supplementary Fig. 2:**
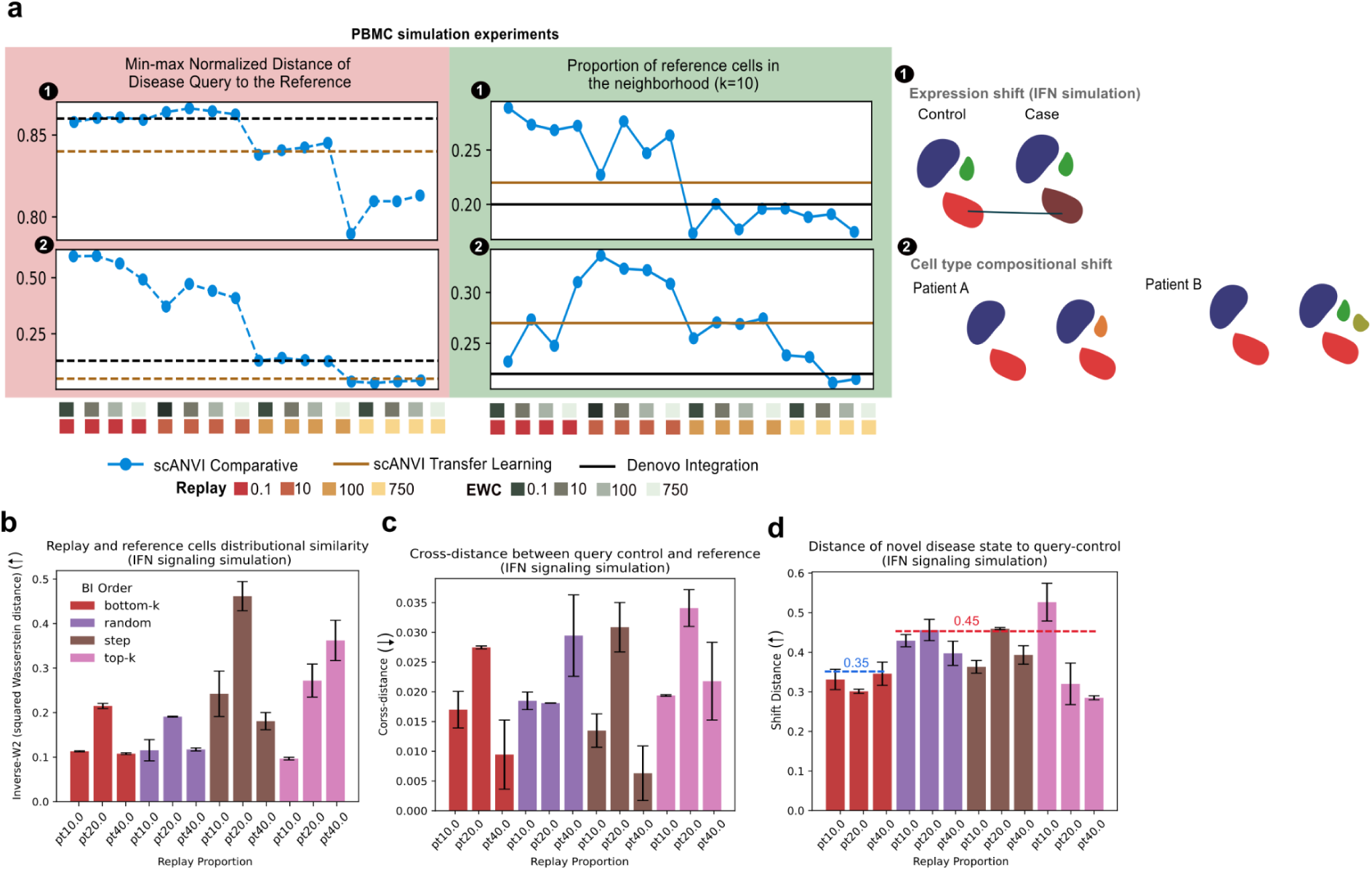
PBMC simulation benchmarks on Replay buffer management for model adaptability.

We found that the distribution defined by the replayed cells is the closest to the reference when the buffer is generated through the “step” approach, indicated by the largest inverse 2-wasserstain squared distance between the replay and reference cells for all buffer sizes compared to random selection, “bottom-k” (**Supplementary Fig.2 b**). The “top-k” scheme was also found to result in a large inverse 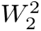 distance for 20% and 40% buffer sizes. This suggests that the “step” and “top-k” approaches can result in minimal distortion of the reference post CL training, hence preservation of biological states in the base cVAE model.

We additionally observed that overall there is a good mixing of cells from query control and reference cells for all buffer management strategies as indicated by small difference (max 0.037) in the distance between query control and reference cells, and average within-query control and within-reference distances, called *cross-distance* (**Supplementary Fig. 2c**).

Since integration of normal and query cells has worked in all runs, we next investigated the preservation of the novel disease state that was simulated in the query-case by measuring the distance of the disease cells to query control only (**Supplementary Fig. 2d**), named as the *Shift Distance*, informed by the fact there is no need to consider cross query-reference distances for normal and control cells, as they mix reasonably well. The results suggest that both random and “step” approaches detect the disease state with a 0.1 difference larger score compared to the “bottom-k” approach. The “top-k” approach at 10% buffer size was also found to make the disease state detectable from the controls, however, the variance of

The “bottom-k” approach selects the most confident cells for replay, “top-k” selects the most uncertain and “step” uses a mix of both confident and cells with high variance uncertainty. Our simulation experiments suggest that the “step” approach introduces smallest distortions to the reference atlas during update by CL, while also resulting in a large distance of the disease state from the control cells in the query. Therefore, we believe that a model trained using CL coupled with a buffer generated through the BI-“step” approach can achieve optimal adaptability.

## Supplementary Note 4: Context-specificity of cell state transitions in response to perturbations

We extended the quantification of state transitions towards Archetypes 1, 3 and 15 from the Epi-CRC atlas to all nine cell lines in Tahoe-100M. We found that the perturbations in LoVo (**Supplementary Fig. 3a**) and RKO (**Supplementary Fig. 3b**) cell lines tend to induce transitions along the malignant and normal cell states. The perturbation in SW1417 (**Supplementary Fig. 3c**) and LS180 (**Supplementary Fig. 3d**) cell lines tend to induce transitions towards a hybrid state of all three normal - absorptive intestine, malignant - ISC-like and pre-malignant - neuroendocrine states, but also predominantly the pre-malignant - neuroendocrine state. SW1417 and LS180 are known to be chemo-tolerant and have secretory/goblet-like phenotype^83–85^, suggesting that their response similarity can be explained by the similarity in context.

In HCT15 and HT-29 (**Supplementary Fig. 3e and f**, respectively), perturbations induced transitions along the normal-neuroendocrine axis, and COLO 205 (**Supplementary Fig. 3g**) and SW48 (**Supplementary Fig. 3h**) are enriched in transitions at the normal state. We found that perturbations can induce transitions to all of the states in SW480 (**Supplementary Fig. 3i**). Overall, we observed that perturbations are more likely to induce transitions along one or several axes, rather than inducing complete transitions to a single state, and that these transitions are strongly context-specific. This finding reveals important insights for the future development of perturbation response models that can generalize between different pre-clinical models and nomination of perturbations for experimental design.

**Supplementary Fig. 3:**
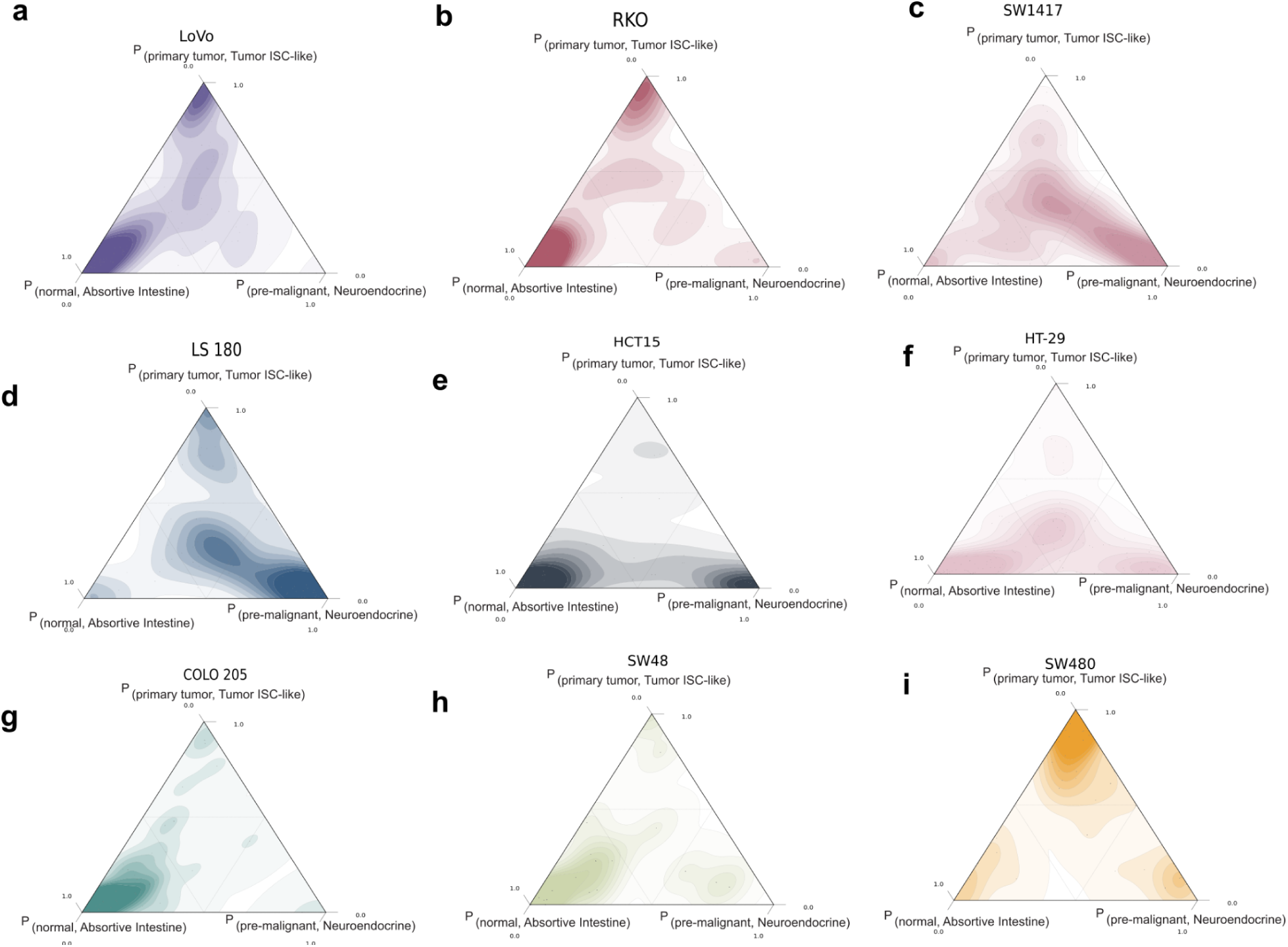
Context-specificity of cell state transitions in response to perturbations.

## Notes

### Summary of Updates

Three of the references were misplaced in version 2. A correction has been applied in the new version.

