## Extended Data Figures for "Perturbation-guided mapping of colorectal cancer cell states to causal mechanisms"

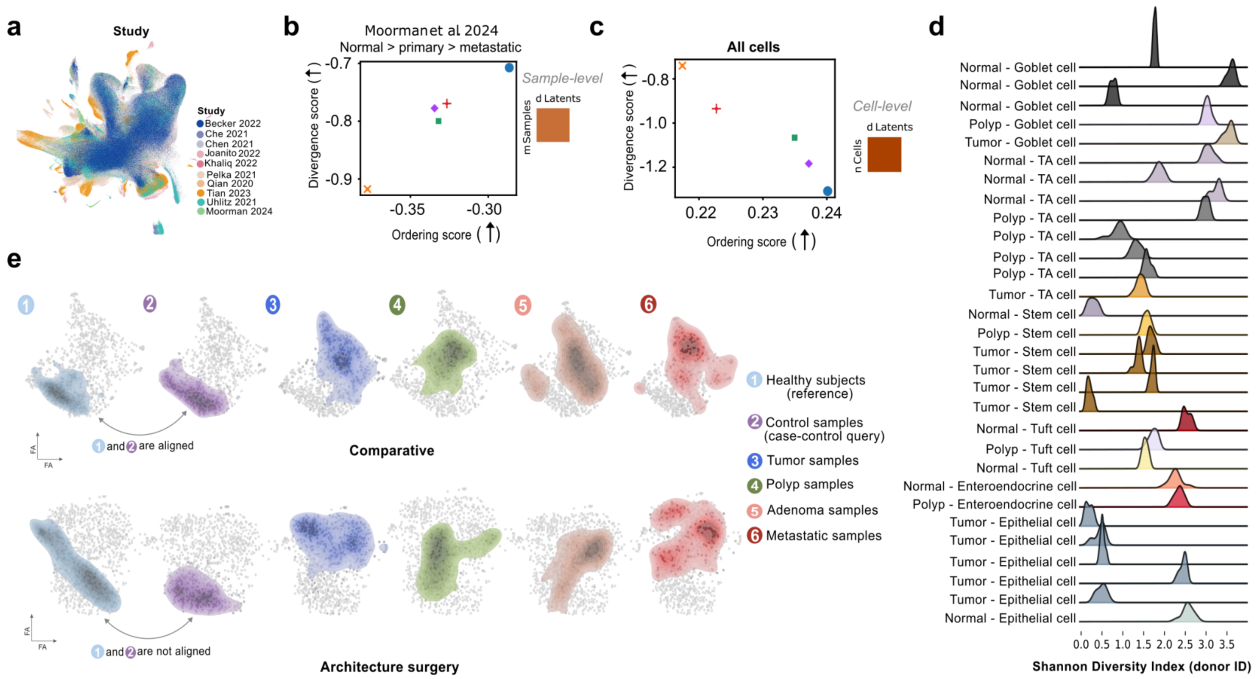


**Extended Data Fig. 1: Preservation of healthy and disease states in the comparative Epi-CRC atlas. (a)** UMAP of integrated Epi-CRC cells colored by study shows study-specific states and good general mixing of cells from different studies**. (b)** Divergence and ordering scores for capturing normal-primary-metastasis transition for the samples in Moorman *et al*. **(c)** Divergence and ordering score for capturing transition to malignancy at the cell level in the Epi-CRC atlas. **(d)** Shannon Diversity Index showing cell mixing by donor ID in a representative group of cell types from normal, polyp and tumour stages. As expected, low mixing is observed in tumor epithelial cells that are patient-specific; high mixing is observed in cell types which are not expected to exhibit patient-specific variation. **(e)** FA plots highlighting healthy samples from the reference, control samples in the query, tumour, polyp, adenoma and metastatic samples. While healthy-reference and query-control samples are distributionally aligned in our comparative approach for continual expansion, expansion via architecture surgery yields distributions of healthy-control samples that are not aligned between reference and query.

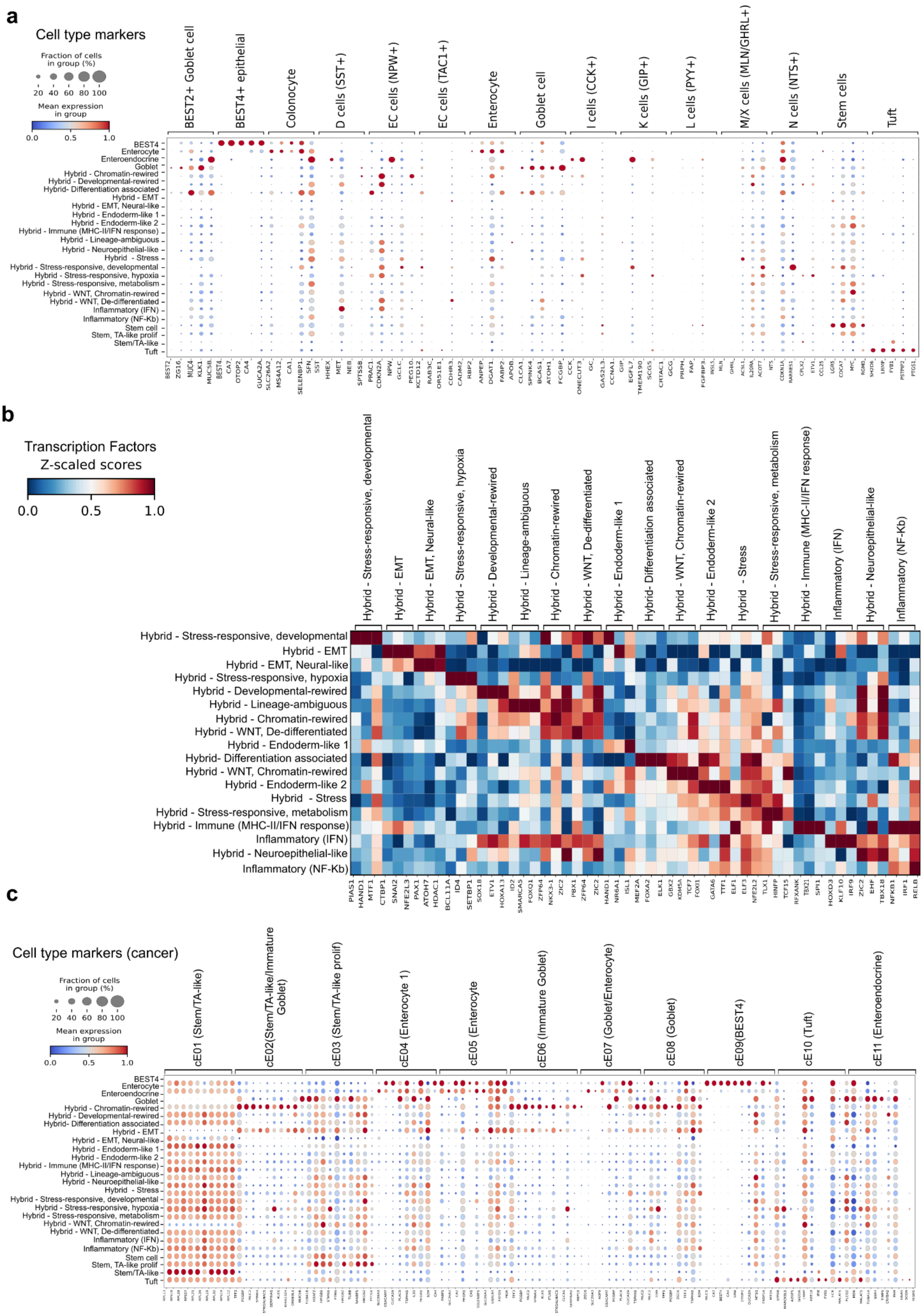

a
Cell type markers
Fraction of cells in group (%)
20 40 60 80 100
Mean expression in group
0.0 0.5 1.0
BEST2+ Goblet cell
BEST4+ epithelial
Colonocyte
D cells (SST+)
EC cells (NPW+)
EC cells (TAC1+)
Enterocyte
Goblet cell
I cells (CCK+)
K cells (GIP+)
L cells (PYY+)
M/X cells (MLN/GHRL+)
N cells (NTS+)
Stem cells
Tuft
BEST4
Enterocyte
Enteroendocrine
Goblet
Hybrid - Chromatin-rewired
Hybrid - Developmental-rewired
Hybrid- Differentiation associated
Hybrid - EMT
Hybrid - EMT, Neural-like
Hybrid - Endoderm-like 1
Hybrid - Endoderm-like 2
Hybrid - Immune (MHC-II/IFN response)
Hybrid - Lineage-ambiguous
Hybrid - Neuroepithelial-like
Hybrid - Stress
Hybrid - Stress-responsive, developmental
Hybrid - Stress-responsive, hypoxia
Hybrid - Stress-responsive, metabolism
Hybrid - WNT, Chromatin-rewired
Hybrid - WNT, De-differentiated
Inflammatory (IFN)
Inflammatory (NF-Kb)
Stem cell
Stem, TA-like prolif
Stem/TA-like
Tuft
b
Transcription Factors
Z-scaled scores
0.0 0.5 1.0
Hybrid - Stress-responsive, developmental
Hybrid - EMT
Hybrid - EMT, Neural-like
Hybrid - Stress-responsive, hypoxia
Hybrid - Developmental-rewired
Hybrid - Lineage-ambiguous
Hybrid - Chromatin-rewired
Hybrid - WNT, De-differentiated
Hybrid - Endoderm-like 1
Hybrid- Differentiation associated
Hybrid - WNT, Chromatin-rewired
Hybrid - Endoderm-like 2
Hybrid - Stress
Hybrid - Stress-responsive, metabolism
Hybrid - Immune (MHC-II/IFN response)
Inflammatory (IFN)
Hybrid - Neuroepithelial-like
Inflammatory (NF-Kb)
PIAS1
HAND1
MTF1
CTBP1
SNAI2
NFE2L3
PAX1
ATOH7
HDAC1
BCL11A
ID4
SETBP1
SOX18
ETV1
HOXA13
ID2
SMARCA5
FOXQ1
ZFP64
NKX3-1
ZIC2
PBX1
ZFP64
ZIC2
HAND1
NR6A1
ISL1
MEF2A
FOXA2
ELK1
GBX2
KDM5A
TCF7
FOXI1
GATA6
TFF1
ELF1
ELF3
NFE2L2
TLX1
HINFP
TCF15
RFXANK
TBX21
SPI1
HOXD3
KLF10
IRF9
ZIC2
EHF
TBX18
NFKB1
IRF1
RELB
c
Cell type markers (cancer)
Fraction of cells in group (%)
20 40 60 80 100
Mean expression in group
0.0 0.5 1.0
cE01 (Stem/TA-like)
cE02(Stem/TA-like/Immature Goblet)
cE03 (Stem/TA-like prolif)
cE04 (Enterocyte 1)
cE05 (Enterocyte
cE06 (Immature Goblet)
cE07 (Goblet/Enterocyte)
cE08 (Goblet)
cE09(BEST4)
cE10 (Tuft)
cE11 (Enteroendocrine)
BEST4
Enterocyte
Enteroendocrine
Goblet
Hybrid - Chromatin-rewired
Hybrid - Developmental-rewired
Hybrid- Differentiation associated
Hybrid - EMT
Hybrid - EMT, Neural-like
Hybrid - Endoderm-like 1
Hybrid - Endoderm-like 2
Hybrid - Immune (MHC-II/IFN response)
Hybrid - Lineage-ambiguous
Hybrid - Neuroepithelial-like
Hybrid - Stress
Hybrid - Stress-responsive, developmental
Hybrid - Stress-responsive, hypoxia
Hybrid - Stress-responsive, metabolism
Hybrid - WNT, Chromatin-rewired
Hybrid - WNT, De-differentiated
Inflammatory (IFN)
Inflammatory (NF-Kb)
Stem cell
Stem, TA-like prolif
Stem/TA-like
Tuft

**Extended Data Fig. 2: Expression of canonical cell type markers in normal and tumour cells used to annotate the cells in Epi-CRC. (a)** Cell type marker expression from Elmentaite *et al*. in malignant cells. **(b)** Activity level of transcription factors used to annotate the hybrid cell types in the Epi-CRC atlas. **(c)** Cell type marker expression from Pelka *et al*. in malignant cells.

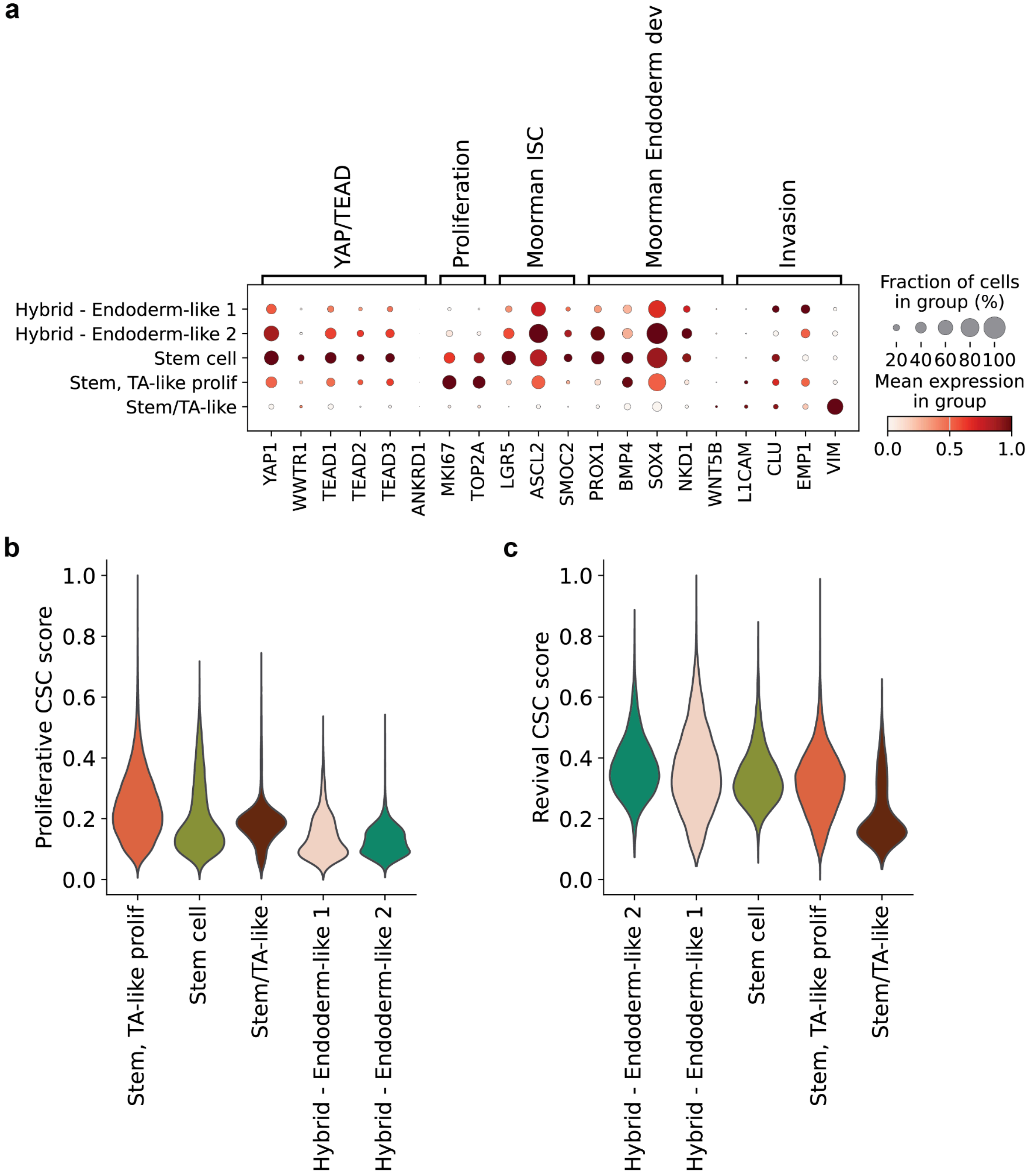

**Extended Data Fig 3: Features of oncofetal cells in hybrid - endoderm-like cell types from the integrated CRC atlas. (a)** Marker gene expression of YAP/TEAD, proliferation, intestinal stem cell (ISC), endoderm development, and invasion by stem cells and endoderm-like states from malignant cells. **(b)** Scaled proliferative cancer stem cell (CSC) scores across the stem cells and endoderm-like state from malignant cells. The cell types and states are ordered by proliferative CSC scores in descending order. **(c)** Similar to **(b)** but for the revival CSC score. The cell types and states are ordered by revival CSC scores in descending order.

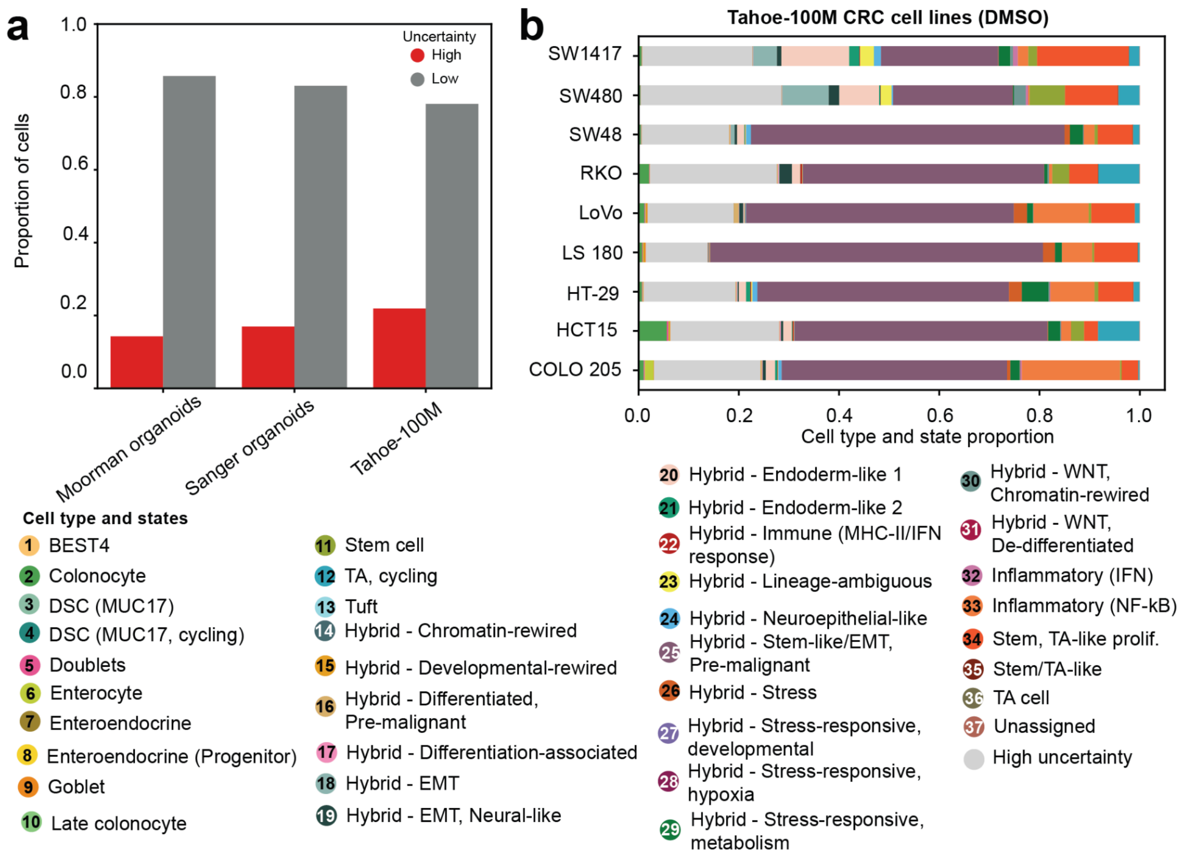


**Extended Data Fig. 4: Proportion of high uncertainty cell labels is highest in Tahoe-100M cell lines. (a)** Comparison of proportion of cells with high (>0.5) and low uncertainty (<0.5) between Tahoe-100M CRC cell lines and CRC organoids**. (b)** Proportion of cell types and states in Tahoe-100M DMSO-treated CRC cell lines. Cells with high uncertainty are coloured grey. A hybrid - stem-like/EMT state associated with pre-malignant samples (state 25) is predominant in cell lines.

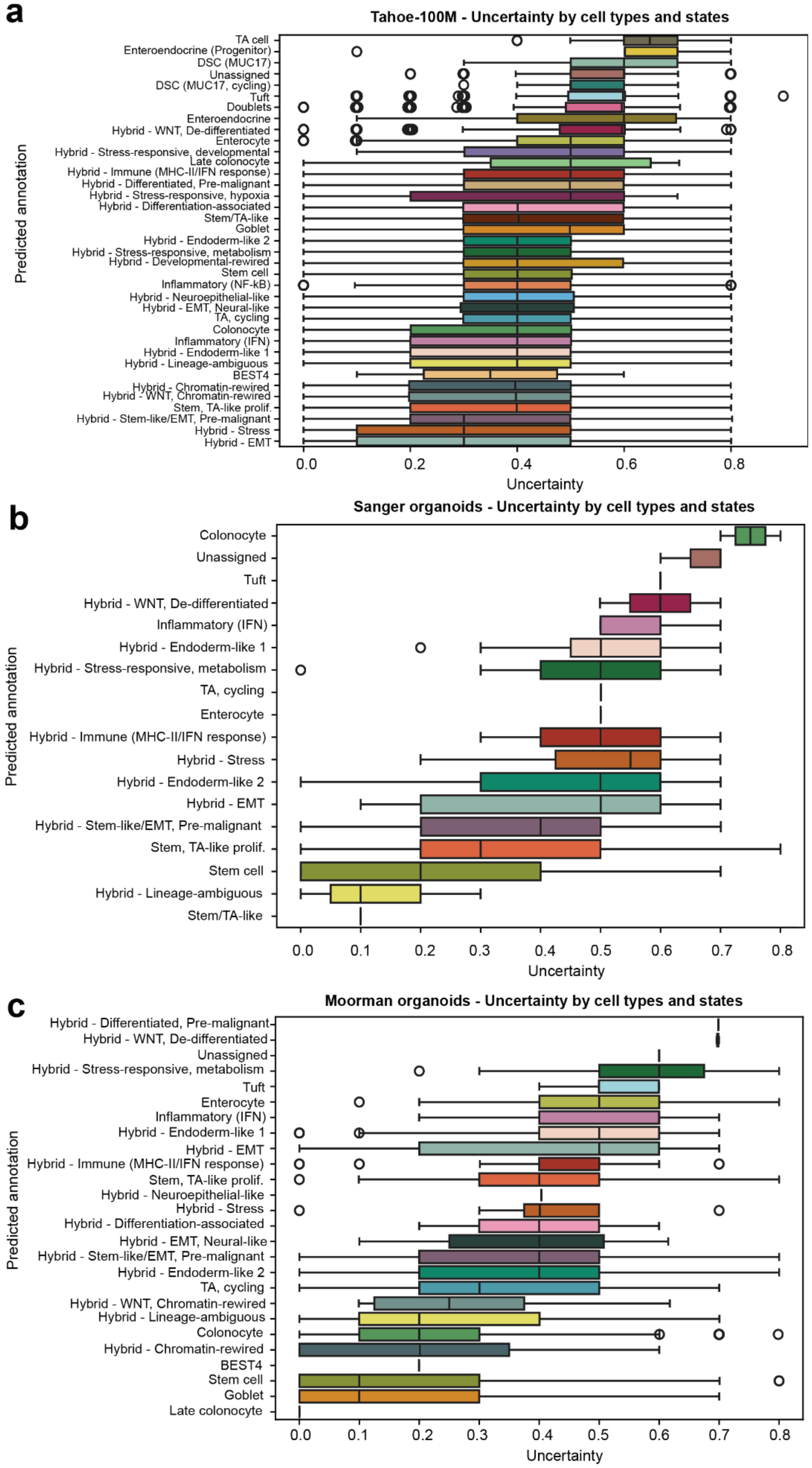


**Extended Data Fig. 5: Distribution of label-transfer uncertainty scores in Tahoe-100M and the CRC organoids. (a)** Distribution of uncertainty scores per cell type in Tahoe-100M, **(b)** Sanger organoids, and **(c)** Moorman organoids.

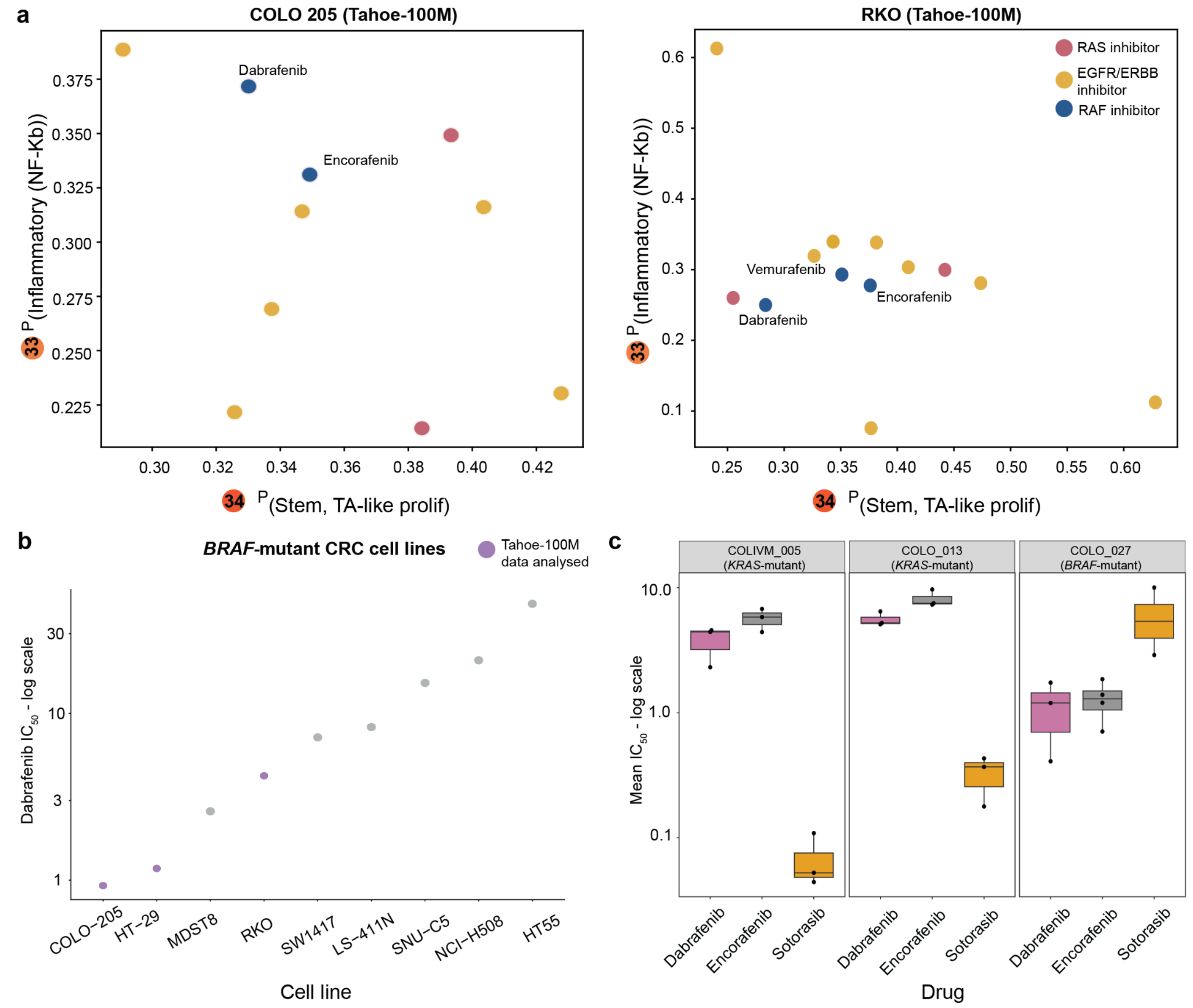


**Extended Data Fig. 6: Contextualizing cell state transitions with drug sensitivity profiles from CRC cell lines and organoids. (a)** Induction of cellular transitions towards the inflammatory (NF-κB) state and stem/TA-like proliferative state by RAS, EGFR/ERBB, and RAF inhibitors in COLO 205 (left) and RKO (right) cell lines. **(b)** Dabrafenib sensitivity of *BRAF*-mutant CRC cell lines, measured by $IC_{50}$ (log scale), from the Genomics of Drug Sensitivity in Cancer 2 (GDSC2) database. The cell lines are ordered by reducing sensitivity towards dabrafenib. Tahoe-100M CRC cell lines are coloured in purple. **(c)** Drug sensitivity profiles of Sanger CRC organoids. $IC_{50}$ values are microMolar. Boxes show the distribution mean $IC_{50}$ (log scale) coloured by drug. Whiskers represent the variance in mean $IC_{50}$ (log scale) across the biological replicates with each point indicating an individual biological replicate. Lower $IC_{50}$ values indicate greater sensitivity towards targeted inhibition.

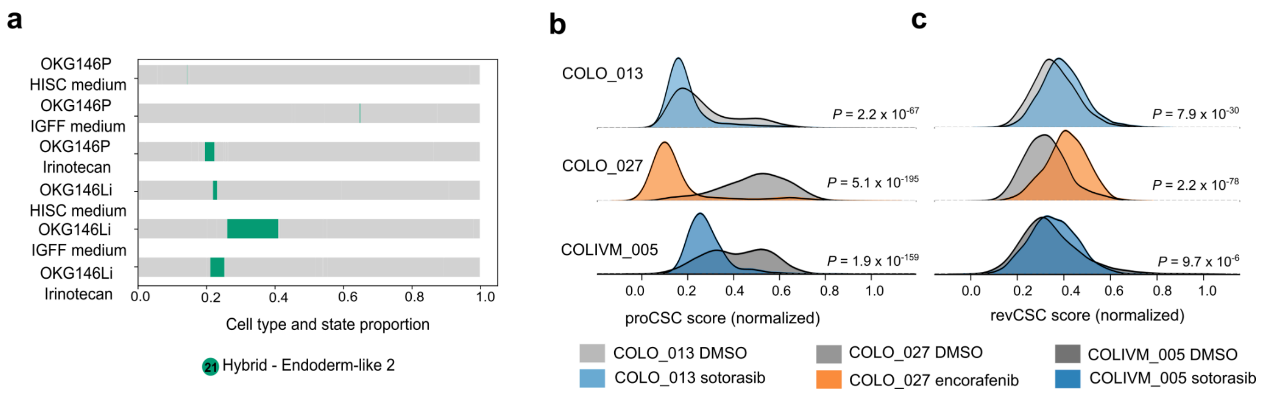


**Extended Data Fig. 7: Orthogonal validation of cell state transitions. (a)** Changes in "Hybrid - Endoderm-like 2" state abundance in *Moorman et al.* organoids pre and post irinotecan treatment. **(b-c)** Normalised expression score for proCSC and revCSC before and after treatment in the Sanger organoids. P-values were calculated using a two-sided Mann-Whitney U test and were adjusted using the Benjamini-Hochberg method.
